# Sediment depth-dependent trait expression controls species contributions to nutrient cycling

**DOI:** 10.64898/2026.09.04.748838

**Authors:** Adam Porter, Martin Solan, Jasmin A. Godbold, Vassilis Kitidis, Mara Fischer, Tara Williams, Callum Roberts, Ceri Lewis

## Abstract

Global biodiversity loss is restructuring ecological communities and compromising ecosystem functioning. Yet, we still lack an empirical understanding of how specific trait combinations regulate ecosystem processes or how their functional expression is organised within the immediate environment. Here, we investigated trait-expression across 17 functionally contrasting sediment-dwelling marine invertebrate species. We show that while sediment reworking and burrow ventilation are depth dependent, their influence on nutrient concentrations are highly nutrient-specific. Our analyses reveal that nutrient concentrations are better explained by continuous, spatially explicit trait combinations than by categorical bioturbation modes. Surface modification was associated with higher ammonium concentrations, moderated by ventilation, while deeper sediment reworking and biomass were associated with higher nitrite concentrations. Phosphate relationships were weak and nitrate was not explained by the measured traits. These results establish a mechanistic baseline for species-specific trait-nutrient relationships under controlled conditions. This shift from categorical trait assignments to a spatially explicit, empirically resolved trait-interaction framework provides a mechanistic foundation for predicting how changes in species composition may alter benthic nutrient cycling.

## Introduction

Understanding of the context-dependent contributions species make to ecosystem process and functioning is critical to the prediction of how intensified levels of human disturbance will alter ecosystems, making it a cornerstone of effective conservation in an era of rapid global change^1,2^. Yet, despite growing evidence that long-term ecosystem persistence depends on sustaining ecological processes^3^, a disproportionate focus on structural biodiversity continues to frustrate progress in this area by failing to capture the subtle but critical degradation of functional processes, particularly at scales that are relevant for conservation decision-making^4–8^. At the community scale, the distribution of functional traits within biological assemblages determines which ecosystem functions are maintained and, by extension, how the community responds to disturbance^9^. Hence, trait-based approaches - analytical frameworks that quantify measurable organismal characteristics (e.g. morphological, physiological, phenological or behavioural traits) and relate them to ecological processes^10^ - can provide mechanistic insights into how species traits mediate responses to environmental conditions. A reoccurring problem, however, is that the expression of traits is not necessarily consistent across different biotic and/or abiotic contexts^11–13^, and communities can show idiosyncratic patterns in response depending on which traits are lost, gained or moderated as well as which processes are considered^14–16^. However, local environmental conditions^17,18^, intraspecific variation^19,20^, compensatory responses^21,22^, arrangement of dominance patterns^23^, and context-dependent interactions^24^ can add substantial complexity and variability into community dynamics, often obscuring observed patterns and limiting the ability of simple categorical metrics^25^ of trait dominance and diversity to fully capture functional relationships^19,26,27^. This means that species with similar roles do not always respond in the same way to the environmental circumstances that are presented, adding credence to the view that direct measurements of ecosystem functions, such as nutrient fluxes, against particular conditions and circumstances will provide a more robust means of quantifying the functional roles of species^13^.

Benthic marine invertebrates display diverse sediment-reworking behaviours that have been grouped into broad functional types (e.g. bioturbation modes^28^) or described using measurable traits^13,29,30^. Bioturbation activities include a suite of behavioural processes (sediment reworking, burrow ventilation) that modify sediment structure, solute transport, and redox gradients, altering microbial activity and, ultimately, nutrient fluxes and sediment biogeochemistry ^31,32^. Trait-based analyses in macrobenthic systems show that functional attributes such as burrowing depth, sediment mixing intensity, and ventilation potential – alone or in combination – can influence nutrient regeneration, organic matter transformation, carbon sequestration, and microbial processing^5,33^. Indeed, species classified within the same bioturbation mode may exert contrasting, or even opposing, effects on different biogeochemical processes, depending on which traits are expressed and at what intensity^34^.

However, species that have been considered functionally equivalent based on their morphology or behaviour^35^ can fall into separate functional groups when their contributions to ecosystem function are measured directly^13^. In addition, intra-specific variability of trait-expression due to differences in body size or burrowing depth^36^ can disproportionately affect mediation of ecosystem functioning above differences in species identity^19^. For example, polychaetes such as *Nephtys hombergii* and *Hediste diversicolor* are often treated as functional equivalents based on particle mixing but can have distinct functional contributions when their effects on ammonium flux are quantified^13^. These findings demonstrate that it is unlikely that any single trait or categorical grouping can adequately capture both ecosystem process and associated ecosystem functioning, reinforcing the need to move beyond trait proxies to direct functional measurement.

These conceptual challenges are particularly significant given the vast spatial scale at which sediment mixing by benthic invertebrates operates. Globally, the mixed depth constitutes at least 13,700 km^3^ −32,500km^3^ of sediment (equivalent to 5.4-16.4 times the volume of Mount Everest)^37^, yet forecasting the functional consequences of altered biodiversity ^14,28^ remains challenging, because individual components of bioturbation do not contribute uniformly across biogeochemical pathways^25^ and sediment reworking behaviours are dynamic, context dependent, and unevenly distributed across taxa^38^. The location and timing of trait expression can also be important; the traits of the species that reside at the sediment–water interface, or that occupy deeper parts of the sediment profile, tend to be disproportionately expressed and are most important for sustaining biogeochemical functioning^39^. As a result, aggregated indices of bioturbation often obscure the mechanistic links between behaviour and function^5,40,41^ and traits that are influential for one function may contribute little to others^4,42,43^, even among closely linked processes. Further, based on morphological and behavioural characteristics, species considered to be functionally equivalent can fall into separate functional groups based on observed contributions to ecosystem functioning^13^. Hence predicting how biodiversity change will affect nutrient cycling requires more than knowing species identities alone; it requires the identification of traits that drive specific functions and how and when they are expressed under particular abiotic and/or biotic circumstances. Specifically, while the effects of individual traits on benthic nutrient cycling are well-documented, we lack an empirical understanding of how contributions of traits expressed by individual species influence nutrient responses through their spatial organisation in the sediment profile^44^. Resolving these relationships requires experimental approaches that control for environmental stochasticity to isolate fundamental trait-function relationships. Although such systems intentionally simplify natural conditions, they enable direct quantification of trait expression and its translation into ecosystem processes, providing the mechanistic baseline necessary to evaluate how these relationships shift across diverse environmental contexts.

Such uncertainties have important implications under anthropogenic disturbance. Changes in infaunal community abundance and composition driven by bottom trawling, invasive species, and pollution have been shown to alter sediment-dwelling communities and shift nutrient cycling dynamics^45–48^. Disturbance can restructure the distribution and intensity of functional traits, such that increases in the biomass or abundance of surviving taxa may compensate for some ecosystem processes but fail to restore others that depend on specific trait combinations or high-intensity expressions. Consequently, alterations in biodiversity can result in persistent functional change even where total biomass or species richness recovers^21,22,49^. While benthic ecosystems support a wide range of ecosystem functions, habitat structuring, and trophic support^50^, we focus here on nutrient cycling as a tractable and mechanistically informative component of ecosystem functioning. Nutrient fluxes integrate complex biological, physical, and microbial interactions, providing a sensitive measure of how faunal traits regulate sediment-water exchange. Crucially, these fluxes govern benthic primary productivity^51,52^, food web stability, and other ecosystem services, such as carbon sequestration^53^. Resolving how individual species and their trait combinations contribute to nutrient generation is therefore essential for developing mechanistic predictions of how ecosystem functioning emerges within sediment communities.

Here, using a functionally diverse set of sediment-dwelling marine invertebrates, we determine species specific contributions to nutrient generation, a process vital to the health of marine ecosystems. Our initial expectation was that species with similar functional characteristics would show similar effects on nutrient generation, and that emergent functional groupings would vary with nutrient identity according to differences in behaviours that alter organism –sediment coupling. Because benthic traits are expressed across different sediment depths, we further expected trait combinations to separate along the vertical sediment profile^39^. We also anticipated considerable intraspecific variability in individual contributions, such that species assigned to distinct categorical groups might overlap functionally depending on the intensity of trait expression.

## Results

### Species-specific differences in sediment mixing traits

Bioturbation and ventilatory activity differed significantly among species (Kruskal–Wallis tests, χ² = 28.1–69.4, all p ≤ 0.05; Fig. 1). We found that *Cerastoderma edule* (bivalve), *Echinocardium cordatum* (heart urchin) and *Priapulus caudatus* (priapulid worm) had the strongest effects on sediment surface properties. Mean surface modification (^SSI^S_mod_; global mean = 5.52 cm^2^ ± 0.89 (s.e.), n = 84; Fig. 1A) generated by the upward transport of particles from depth or the overturning of surface sediments ranged from 0.483 ± 0.183 cm^2^ by *Capitellidae* sp. (polychaete worm) to 21.611 ± 3.182 cm^2^ by *E. cordatum* and mean (± s.e.) surface boundary roughness (^f-SPI^SBR; global mean = 0.61 cm ± 0.04 (s.e.), n = 84; Fig. 1B), ranged from 0.240 ± 0.028 cm by *Notomastus latericeus* (polychaete worm) to 1.240 ± 0.167 cm by *P. caudatus* (Fig. 1b).

**Figure 1.**
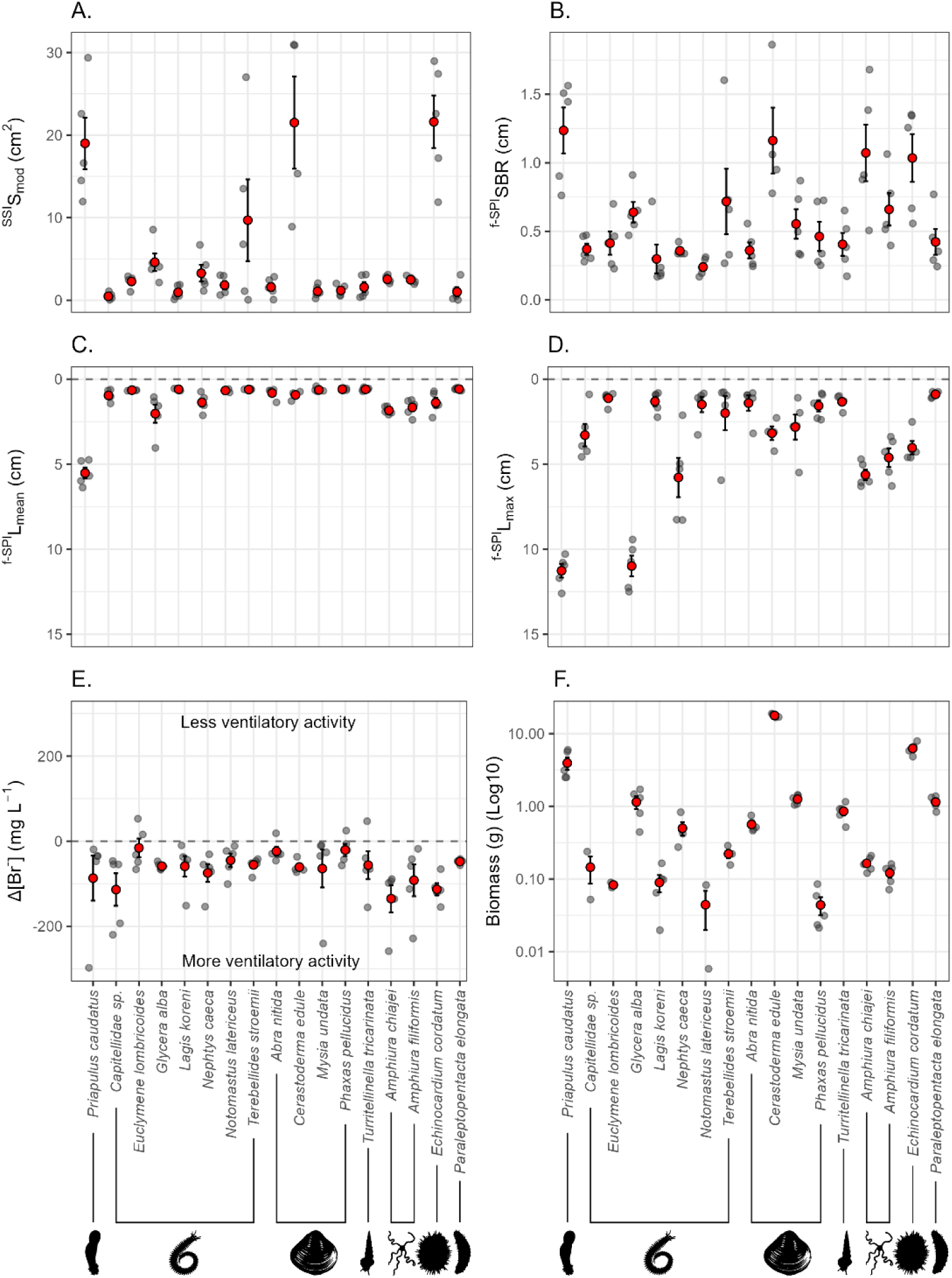
Summary of the species-specific contributions to different components of bioturbation behaviour (ecosystem process) and biomass for representative sediment-dwelling marine invertebrates. Data are shown for (A) surface modification (^SSI^S_mod_), (B) surface boundary roughness (^f-SPI^SBR), (C) mean depth of particle reworking (^f-SPI^L_mean_), (D) maximum depth of particle reworking (^f-SPI^L_max_), (E) ventilatory activity (Δ[Br^−^]) and (F) species biomass (log₁₀-transformed biomass (g)). In each panel, red circles represent the mean values for each species and associated standard error bars (n ≈ 5 individual species^−1^, grey circles). In (F) the y-axis is Log10 transformed to improve visualisation. The silhouette insets indicate broad faunal groups (from left to right: the priapulids (Priapulidae); the polychaetes (Nereididae); the molluscs (Bivalvia and Gastropoda); and the echinoderms (Echinoidea and Holothuroidea)) and are from PhyloPic (https://www.phylopic.org/; Keesey, 2023^54^).

Our results show that mean sediment reworking depth (^f-SPI^L_mean_; global mean = 1.26 cm ± 0.14 (± s.e.), n = 84; Fig. 1C) among species, ranged between 0.585 ± 0.028 cm and 2.03 ± 0.54 cm for all species except *P. caudatus* (^f-SPI^L_mean_, 5.52 ± 0.32 cm). There was large species-specific variation in the maximum mixing depth (^f-SPI^L_max_; 10.41 cm difference; min = 0.89 ± 0.08 cm, max = 11.26 ± 0.4 cm, global mean = 3.69 cm ± 0.36 (± s.e.), n = 84; Fig. 1D). Deepest mixing (^f-SPI^L_max_) occurred in the presence of *P. caudatus* (mean (± s.e.), 11.26 ± 0.4 cm), followed by *Glycera alba* (polychaete worm) (10.98 ± 0.6 cm), *Nephtys caeca* (polychaete worm) (5.78 ± 1.16 cm) and *Amphiura chiajei* (brittlestar) (5.62 ± 0.32 cm). Burrow ventilation (Δ[Br^−^], global mean = −66.1 ± 6.90 mg L^−1^ (± s.e.), n = 84; Fig. 1E) was similar among most species and activity was highest for *A. chiajei*, followed by *Capitellidae* sp., *E. cordatum*, and *Amphiura filiformis* (brittlestar) (Fig. 1E). Overall, individual biomass (global mean = 2.1 g wet weight ± 0.50 (± s.e.), n = 72; Fig. 1F) varied strongly among species ranging from 0.043 ± 0.012 g for *Phaxas pellucidus* (bivalve) to 17.723 ± 0.54 g for *C. edule* (Fig. 1F).

### Relative importance of intraspecific variation in trait expression

Intraspecific variability differed markedly among measured ecosystem processes (bioturbation metrics and ventilatory activity) and among species biomass which scales effect strength. While effect-related metrics such as ventilatory activity (Δ[Br^−^]), surface boundary roughness, and surface modification exhibited high within-species variability (72.3%, 39.2%, and 27.5% respectively), the trait metrics underpinning sediment reworking; maximum and mean reworking depth and biomass showed low intraspecific variability (15.76%, 10.89%, and 1.67%), indicating strong within-species constraint on these traits.

### Species-specific contributions to nutrient concentrations

Ammonium Log-Normal Response Ratio (LNRR; a proportional, log-transformed effect size that stabilises variance and puts increases/decreases on a symmetric scale relative to the no-fauna control) generally indicated greater concentration in the water column than no macrofauna controls across most species. The highest concentrations were observed in *C. edule* (1.457 ± 0.039), *N. latericeus* (1.435 ± 0.181), and *Euclymene lombricoides* (polychaete worm) (1.149 ± 0.304). Notably, only *Paraleptopentacta elongata* (sea cucumber) (−0.062 ± 0.112 LNRR) and *A. nitida* (−0.044 ± 0.173) had lower concentrations than the no macrofauna controls (Fig. 2A).

**Figure 2.**
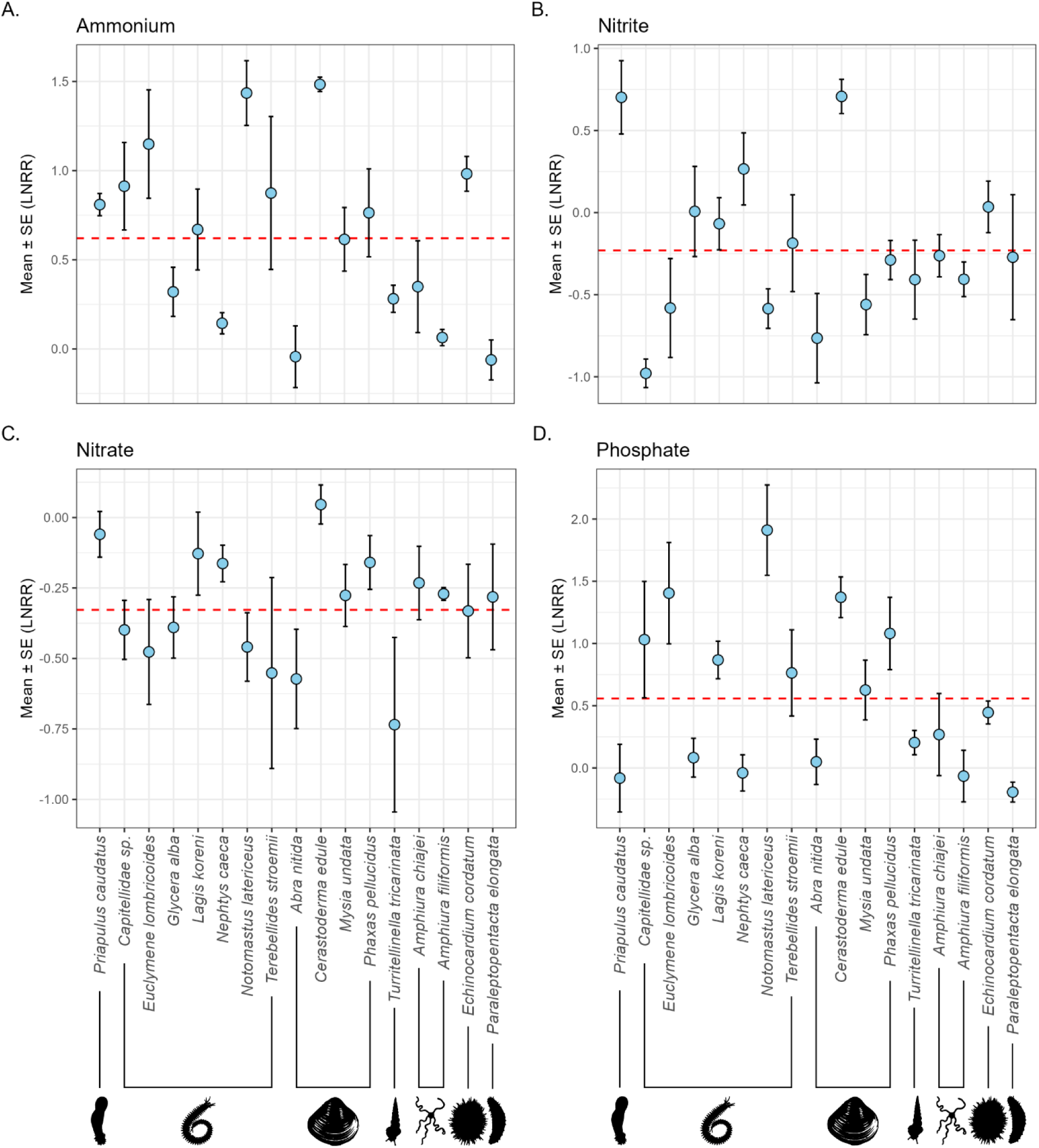
Species-specific nutrient concentrations expressed as log-normal response ratios (LNRR). Panels show mean (± standard error (s.e.)) species contributions to (A) ammonium (NH₄⁺-N), (B) nitrite (NO₂⁻-N), (C) nitrate (NO₃⁻-N), and (D) phosphate (PO₄³⁻-P). The dashed red line indicates the global mean (LNRR) for each nutrient, providing a reference against which species-specific deviations can be assessed (n = 5 for all species except for *C. edule* with *n* = 4).

Nitrite concentration(LNRR) was weakest in the presence of *Capitellidae sp.* and *A. nitida* (Nitrite LNRR −0.979 ± 0.087 and −0.764 ± 0.272 respectively), while the greatest concentrations were observed in treatments containing *P. caudatus* (0.702 ± 0.223) and *C. edule* (0.605 ± 0.492) (Fig. 2B).

Across all species there was evidence of nitrate depletion; it was strongest in the presence of *Turritellinella tricarinata* (auger shell) (−0.735 ± 0.309 LNRR), *A. nitida* (−0.573 ± 0.176 LNRR) and *Terebellides stroemii* (polychaete worm) (−0.552 ± 0.339 LNRR) (Fig. 2C).

Phosphate values (LNRR) indicated predominantly increased concentrations across the species pool, with the strongest responses observed in *N. latericeus* (1.911 ± 0.363), *E. lombricoides* (1.405 ± 0.408), and *C. edule* (1.270 ± 0.165). Elevated phosphate release was also observed in *P. pellucidus* (1.081 ± 0.290), *Capitellidae* sp. (1.031 ± 0.467), and *Lagis koreni* (polychaete worm) (0.868 ± 0.151). In contrast, low or near-neutral responses were recorded in *G. alba* (0.083 ± 0.156), *A. nitida* (0.049 ± 0.182), *N. caeca* (−0.039 ± 0.146), *A. filiformis* (−0.065 ± 0.207), *P. caudatus* (−0.082 ± 0.271), and *P. elongata* (−0.194 ± 0.080) (Fig. 2D).

### Trait combinations predict species-specific nutrient responses

Using principal component analysis and Pearson’s correlations (Supplementary Fig. S1 and S2) we identified surface modification (^SSI^S_mod_), maximum sediment reworking depth (^f-SPI^L_max_) and ventilatory activity (Δ[Br^−^]) as the strongest variables to represent the different components of ecosystem processes (deep burrowing, upward transport and surficial overturning of particles and burrow ventilation, respectively).

### Surface modification and ventilatory activity interactively regulate ammonium concentrations

Ammonium (NH₄⁺–N) concentration was driven by organism size, surface modification and ventilatory activity. Likelihood-ratio test identified NH₄⁺–N concentration to be driven by the interaction between surface modification and ventilatory activity (*Χ*² = 6.38, d.f. = 1, p = 0.012), with the interaction model also favoured over the corresponding additive model by AIC (ΔAIC = 4.38), and independently by biomass (*Χ*² = 6.10, d.f. = 1, p = 0.013). Overall surface modification showed the strongest statistical support among fixed effects (*Χ*² = 12.96, d.f. = 1, p < 0.001), followed by biomass (*Χ*² = 6.10, d.f. = 1, p = 0.013) and ventilatory activity (*Χ*² = 5.71, d.f. = 1, p = 0.017). Fixed effects (marginal R^2^) explained 37.0% of the variance, increasing to 53.4% when species-level random effects were included (conditional R^2^). Specifically, we observed a negative effect of surface modification and ventilatory activity on NH₄⁺–N (*t*(71.14) = −2.61, *p* = 0.011; Fig. 3A); while surface disturbance generally stimulated elevated ammonium concentration, this effect was offset by higher levels of ventilation. Hence, our findings reveal that while both sediment disturbance and ventilation contribute to ammonium dynamics, their combined influence is non-additive. This reflects a complex interaction between physical reworking and advective porewater exchange, where one process moderates the functional outcome of the other.

**Figure 3.**
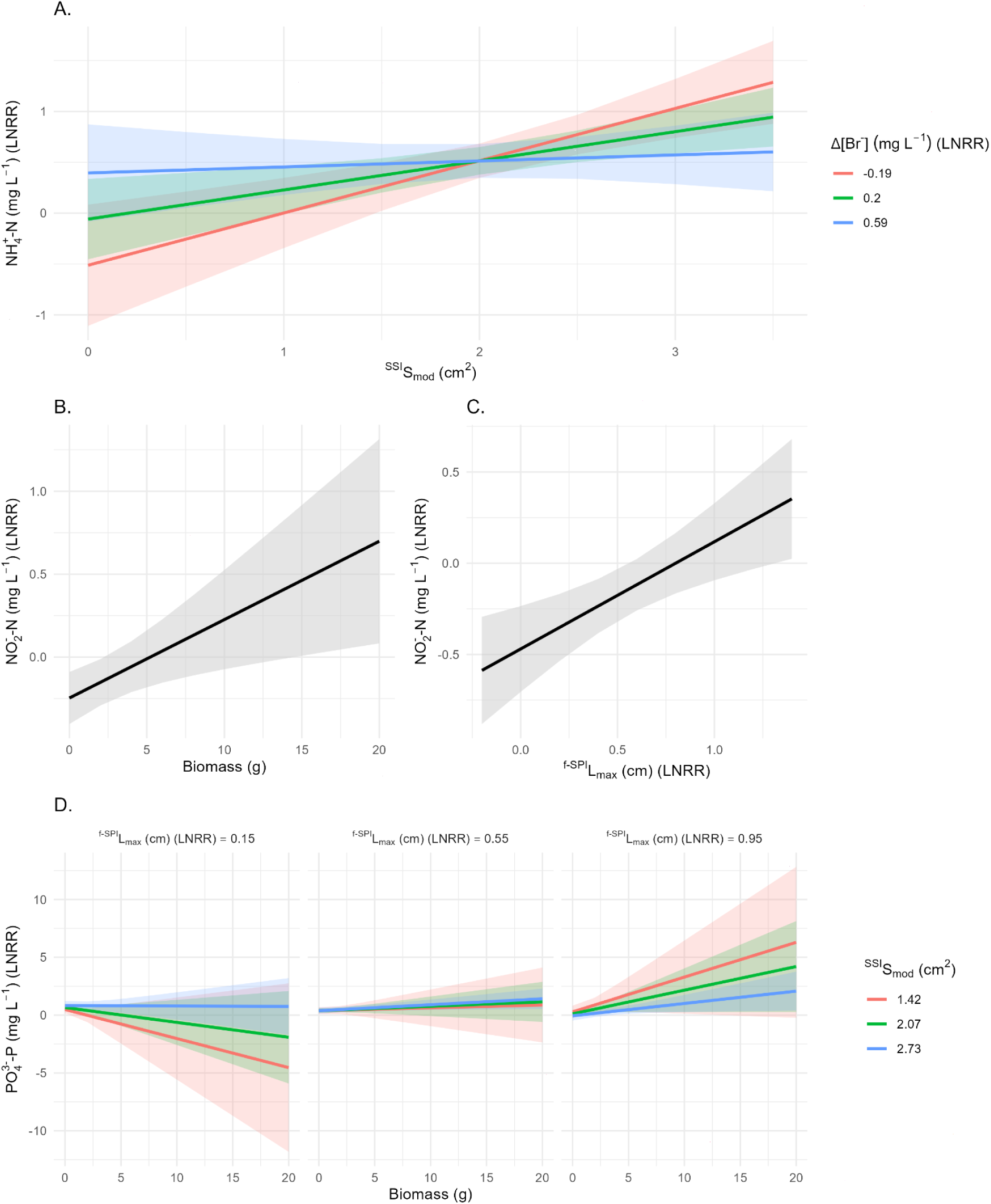
Trait–nutrient relationships derived from linear mixed-effects models. Panels show the significant model-predicted marginal effects of biomass, maximum sediment reworking depth (^f-SPI^L_max_), surface modification (^SSI^S_mod_), and ventilatory activity (Δ[Br⁻]) on nutrient concentrations expressed as log response ratios (LNRR). (A) Ammonium (NH₄⁺-N) responses to surface modification (^SSI^S_mod_) across levels of ventilatory activity, illustrating an interaction between surface processes and ventilation. (B-C) Nitrite (NO₂⁻-N) responses to biomass (B) and maximum sediment reworking depth (^f-SPI^L_max_) (C). (D) Phosphate (PO₄³⁻-P) responses to biomass across gradients of surface modification (^SSI^S_mod_) and maximum sediment reworking depth (^f-SPI^L_max_), reflecting a weak and non-significant interaction between surface modification and depth, with biomass scaling the magnitude of responses. Solid lines represent model predictions with shaded areas indicating 95% confidence intervals. Colours and facets show model predictions evaluated at trait values set to the mean and ±1 standard deviation (−1 SD, Mean, +1 SD), illustrating how nutrient concentrations depend on trait interactions rather than single predictors. Nitrate (NO₃⁻-N) showed no significant response to our measured variables and so is not plotted.

Mean (± s.e.) ammonium concentration was highest in the presence of shallow-mixing, high-biomass surface modifiers (e.g., *C. edule*, ^SSI^S_mod_ = 21.51 ± 5.58 cm², NH₄⁺–N (LNRR) = 1.45 ± 0.04; *E. cordatum*, ^SSI^S_mod_ = 21.61 ± 3.18 cm², NH₄⁺–N (LNRR) = 0.98 ± 0.1; Fig 1A, 2A, and 3C), consistent with the positive main effects of biomass and surface modification identified in the model; however, the key interaction was between surface modification and ventilation, with ventilation dampening the effect of surface disturbance on ammonium release., and reduced in the presence of deeper mixing species (e.g., *N. caeca*, ^f-SPI^L_max_ = 5.78 ± 1.16 cm, NH₄⁺–N (LNRR) = 0.14 ± 0.06; *G. alba*, ^f-SPI^L_max_ = 10.98 ± 0.60 cm, NH₄⁺–N (LNRR) = 0.32 ± 0.14) (Fig. 1D, 2A, and 3B). Strong ventilatory activity by brittle stars (*A. filiformis*, Δ[Br⁻] = −91.87 ± 37.55, NH₄⁺–N (LNRR) = 0.06 ± 0.05; *A. chiajei*, Δ[Br⁻] = −135.08 ± 31.98, NH₄⁺–N (LNRR) = 0.35 ± 0.26 (mean ± s.e.)) was associated with lower ammonium concentration despite surface disturbance.

The contribution of individual species to unexplained variance differed markedly, indicating heterogeneity in how closely species conformed to trait-based expectations. Species such as *N. latericeus*, *Abra nitida* (glossy furrow shell), and *A. filiformis* contributed disproportionately to the random-effects variance (Supplementary Fig. S3C), despite exhibiting only moderate surface modification values (1.894 ± 0.501, 1.604 ± 0.491, and 2.485 ± 0.183 cm², respectively; mean ± s.e.). In contrast, species such as *E. cordatum* (21.611 ± 3.182 cm²) and *P. caudatus* (19.0 ± 3.13 cm²) exhibited high and consistent levels of surface modification but contributed relatively little unexplained variance, indicating that their effects on ammonium concentrations were well captured by the fixed effects.

Consistent with these species-level patterns, *C. edule* (NH₄⁺–N LNRR = 1.46 ± 0.04), *N. latericeus* (1.44 ± 0.18), and *E. lombricoides* (1.15 ± 0.30) exhibited ammonium concentration values ≥1 SD above the global mean, whereas *A. nitida* (−0.04 ± 0.17) and *P. elongata* (−0.06 ± 0.11) fell below the mean (Figs. 2C, 4).

**Figure 4.**
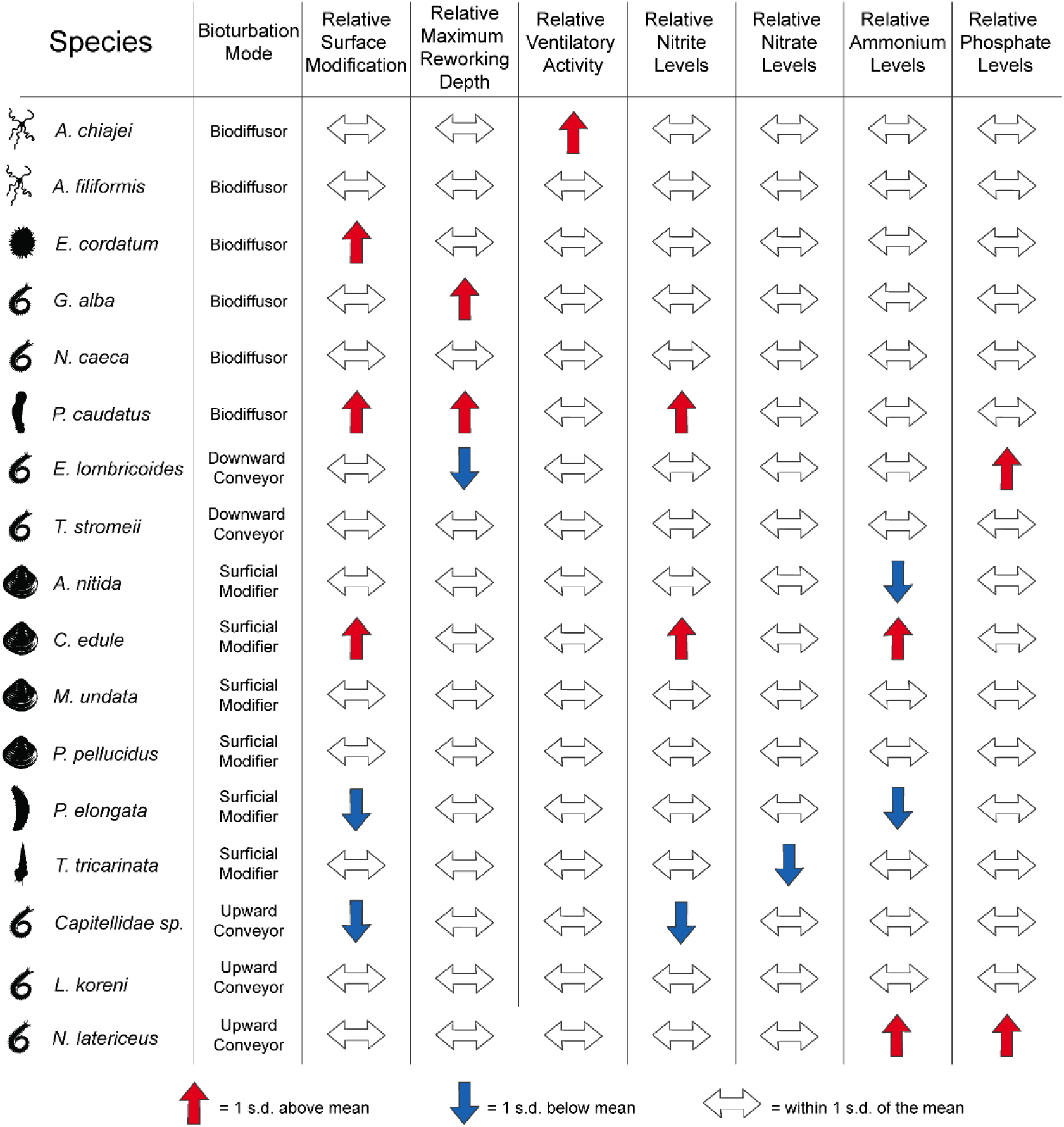
Summary of species-level traits and nutrient concentrations highlighting comparatively higher and lower species-specific contributions to function under experimental conditions. Bioturbation modes (Biodiffusor, Conveyor, Surficial Modifier; modified from Queiros et. al.^30^) are assigned to illustrate the limitations of these categorical designations, as trait-based responses measured here do not map neatly onto them. Relative trait values (surface modification (^SSI^S_mod_), maximum sediment reworking depth (^f-SPI^L_max_), ventilatory activity (Δ[Br⁻])) and nutrient concentrations (ammonium, nitrite, nitrate, phosphate) are shown as deviations from the mean. Red upward arrows indicate values ≥1 standard deviation above the mean meaning more mixing activity, ventilation or greater concentrations, blue downward arrows indicate values ≤1 standard deviation below the mean meaning less mixing, ventilation, or lower concentrations, and grey horizontal arrows indicate values within 1 standard deviation of the mean. This representation distinguishes species that show disproportionally high (consistent positive or negative deviations) or low (responses close to or below the global mean) contributions to functioning within a given set of circumstances.

### Nitrite concentration depends on organism size and sediment reworking depth

In contrast to the dynamics observed for ammonium concentrations, nitrite was driven by the independent and additive effects of organism size and deeper sediment reworking (biomass: *Χ*² = 6.71, d.f. = 1, p = 0.010; sediment reworking depth: *Χ*² = 9.87, d.f. = 1, p = 0.002). Nitrite concentrations increased with both organism size (b = 0.047, SE = 0.017, t(18.08) = 2.81, p = 0.012; Fig. 3B) and maximum sediment reworking depth (b = 0.588, SE = 0.174, t(24.18) = 3.38, p = 0.002; Fig. 3C); the absence of an interaction between these traits, suggests that their species specific contributions to nitrite release operate through distinct pathways, rather than complex trait interdependencies.

High-biomass taxa, such as *C. edule* (17.72 ± 0.054 g; Fig. 1F and 4), exhibited comparatively high nitrite concentrations, consistent with the positive effect of organism size. Species characterised by deeper sediment reworking, including *P. caudatus* (^f-SPI^L_max_ = 11.26 ± 0.40 cm) and *N. caeca* (^f-SPI^L_max_ = 5.78 ± 1.16 cm) aligned with the positive influence of sediment mixing depth. However, deep-burrowing behaviour alone was not a consistent predictor of nitrite flux, as not all deep-dwelling species (e.g. *G. alba*) exhibited strong effects on nitrite concentrations despite deep burrowing (^f-SPI^L_max_ = 10.98 ± 0.60 cm), likely because lower biomass constrained the expression of depth-related effects (Fig. 3A; Figs. 1D, 4).

The fixed effects in the nitrite model (marginal R²) explained 31.6% of the variance in nitrite concentrations, increasing to 39.7% when species-specific random effects were included (conditional R²). This indicates that whilst variation in nitrite concentration is largely captured by additive effects of organism size and sediment reworking depth, but that additional variability is attributable to species-specific differences. Our analyses reveal that several species deviate from model predictions; *Lagis koreni*, *Nephtys caeca*, and *Priapulus caudatus* exhibited higher nitrite concentrations than expected based on their trait values, while *N. latericeus* and *E. lombricoides* closely followed trait-based expectations and contributed little additional unexplained variance (Supplementary Fig. S3A).

### Nitrate concentration is not explained by measured traits

Nitrate (NO₃⁻–N) concentrations were not explained by biomass or measured ecosystem process traits. The best-supported model included only an intercept and a species-level random effect, indicating that variation in nitrate concentration was not associated with the trait-based mechanisms considered here (likelihood-ratio test: *Χ*² = 0.33, d.f. = 1, p = 0.563).

### Surface phosphate concentration is constrained by deep sediment mixing

Phosphate (PO₄³⁻–P) concentrations exhibited weak but complex relationships with organism traits. Model selection did not identify any significant fixed effects (all p > 0.05), indicating limited statistical support for trait-based predictors. However, patterns in the data suggested a negative interaction between surface modification and maximum sediment reworking depth (^SSI^S_mod_ × ^f-SPI^L_max_ : b = −0.648, SE = 0.410, t(67.09) = −1.58, p = 0.119; Fig. 3E), such that increases in phosphate concentrations associated with surface disturbance may be reduced at greater sediment mixing depths, consistent with potential downward transport and retention of phosphate away from the sedimentwater interface. This pattern suggests that increases in phosphate concentrations associated with surface disturbance may be reduced at greater sediment mixing depths, consistent with potential downward transport and retention of phosphate away from the sediment–water interface.

Although our results revealed that trait-based relationships with phosphate concentration were not strong, patterns in the data indicated that at the species level, phosphate concentrations by surface-modifying, shallow-mixing taxa such as *N. latericeus* (mean ± s.e. LNRR = 1.91 ± 0.36), *E. lombricoides* (1.41 ± 0.41), and *C. edule* (1.27 ± 0.17), was well above the global mean (Fig. 2D) and contributed disproportionately to phosphate release (Fig. 4). In contrast, deep mixing taxa such as *G. alba* (^f-SPI^L_max_ = 10.98 ± 0.60 cm) and *P. caudatus* (11.26 ± 0.40 cm) resulted in low or negative phosphate concentrations (0.08 ± 0.16 and −0.08 ± 0.27, respectively; Fig. 2D), consistent with the potential burial and retention of phosphate under deeper sediment reworking.

Fixed effects explained 25.0% of the variance in phosphate concentration (marginal R^2^), increasing to 36.6% when species-level variation was included (conditional R^2^), indicating that a substantial proportion of variability remained unexplained by measured traits. Variance partitioning of random effects revealed that several species deviated from model predictions. Species such as *P. pellucidus*, *T. stroemii*, *N. latericeus*, and *L. koreni* exhibited higher phosphate concentrations than expected based on their trait values, whereas *N. caeca*, *A. nitida*, and *A. filiformis* showed lower-than-expected responses (Supplementary Fig. S3D). Collectively, these results indicate that phosphate concentration exhibits greater interspecific variability relative to the other nutrient pathways we have considered. In this instance, trait-based models capture broad functional trends rather than the nuances of species-specific responses.

### Sensitivity analysis

Although nutrient responses were measured for five individuals per species, except C. edule (n = 4), complete-case replication in the ammonium, nitrite and phosphate models was reduced to n = 2 for Capitellidae sp., Euclymene lombricoides and Notomastus latericeus. Excluding these species produced closely comparable effect estimates: biomass, surface modification and the surface modification × ventilation interaction remained supported for ammonium; nitrite estimates were virtually unchanged; and phosphate relationships remained insignificant. The contribution of ventilation as a main effect in the ammonium model weakened however (p = 0.064; Supplementary Table S2).

### Species-level variance and trait-based predictability

The variance components for the three significant models (nitrite, ammonium, and phosphate) were assessed to understand the variability in the random effect of species identity using Intraclass Correlation Coefficient (ICC). The ICC was 11.7% for the nitrite model, 26.1% for the ammonium model, and 15.4% for the phosphate model, indicating that a larger proportion of unexplained variance in ammonium and phosphate concentrations is structured at the species level compared to nitrite. Species-level residuals from each model, highlighting systematic deviations that remain after accounting for trait effects, are shown in Supplementary Fig. S4.

## Discussion

Our findings demonstrate that species-specific contributions to nutrient concentrations depend not only on the traits a species possesses, but on the location and intensity of their expression within the sediment. A species’ contribution to nutrient concentrations is therefore not a fixed property that can be inferred from trait identity alone. Here, by standardising environmental conditions (individuals from one region, sampling period and incubated in a single sediment type, at 12 °C and under a fixed light regime) were able to establish a mechanistic baseline for species-level trait-nutrient relationships across a diverse pool of benthic invertebrates. We show that spatial structuring of sediment (through surface modification and reworking depth) interacts with ventilation as a regulatory process, with biomass acting to scale the magnitude of effects. Importantly, species-specific contributions to nutrient concentrations were not consistently associated with any single dominant trait but instead reflected additive and interactive combinations of traits expressed across spatial and behavioural dimensions. This framework effectively identifies species whose contributions to functioning are disproportionately high or low relative to their categorical assignments (Fig. 4). While these results establish a mechanistic baseline under controlled conditions, we anticipate that environmental context, species interactions, and sediment properties may moderate the specific parameterization of these relationships. Because the experiment examined isolated individuals, whether the identified relationships combine additively or are modified by facilitation, competition or other interactions within multispecies assemblages remains to be tested directly. Nevertheless, the consistency of the identified relationships provides an empirically resolved, spatially explicit trait framework for generating testable predictions of how changes in assemblage composition may alter benthic nutrient cycling. Conceptually, these nutrient-specific interaction structures illustrate how the location and intensity of trait expression are associated with distinct nutrient responses (Fig. 5A); nitrite increases with both biomass and sediment reworking depth, nitrate remains weakly linked to trait variation (Fig. 5B), ammonium is associated with surface-mediated processes moderated by ventilation (Fig. 5C), and phosphate shows weak and variable relationships with measured traits, suggesting a balance between surface modification and sediment mixing (Fig. 5D).

**Figure 5.**
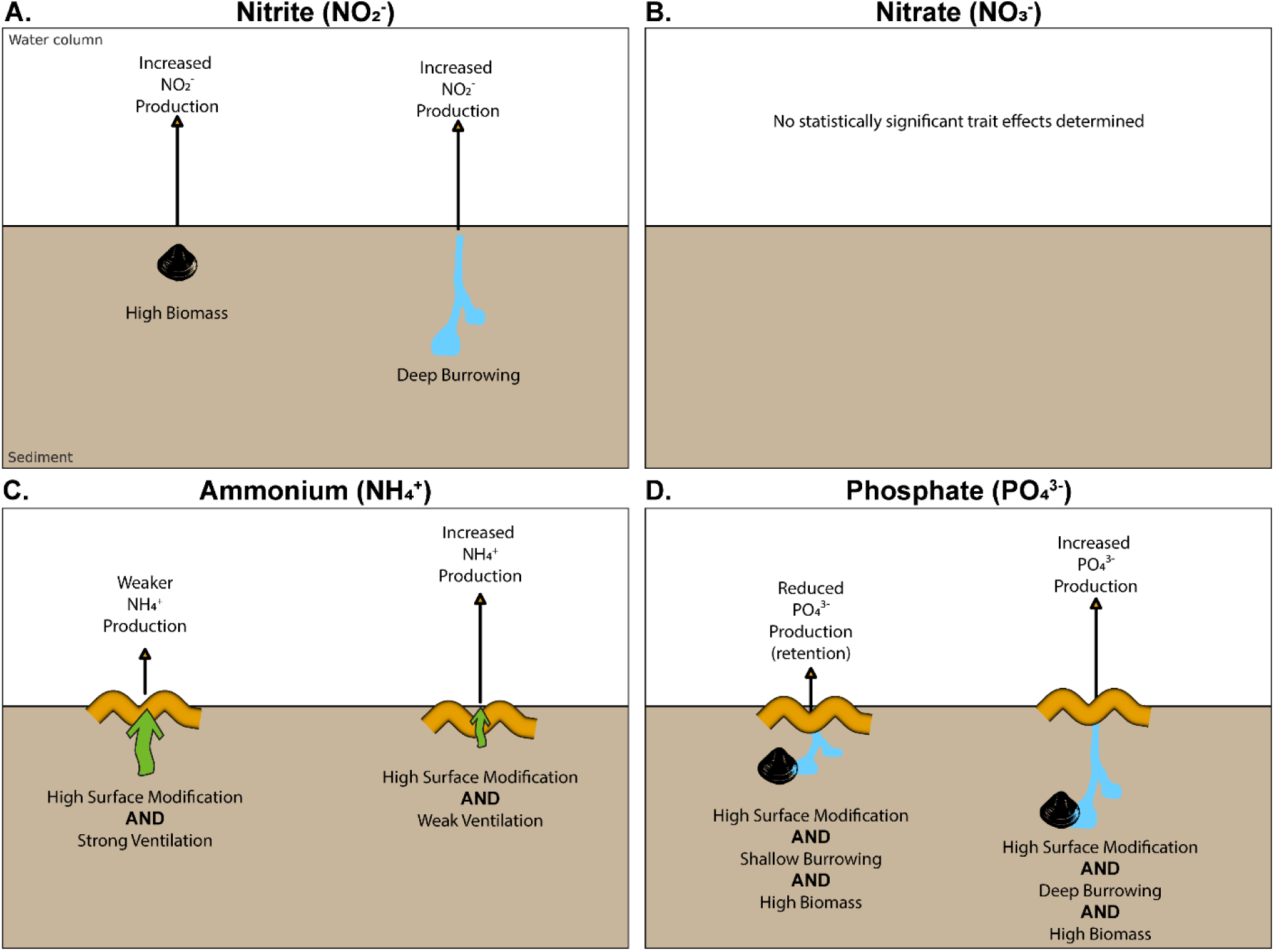
Conceptual synthesis of species-level trait-nutrient relationships observed showing how combinations of biomass, sediment reworking depth, surface modification, and ventilation influence nutrient concentrations. (A) Nitrite (NO₂⁻) concentrations increase with both high biomass and deep burrowing. (B) Nitrate (NO₃⁻) concentration showed limited variability, but there is no evidence that any of the specific traits tested here are influential. (C) Ammonium (NH₄⁺) concentration is driven by surface modification with outcomes regulated by ventilation: strong ventilation reduces elevated ammonium concentrations, whereas weak ventilation allows accumulation and export. (D) Phosphate (PO₄³⁻) concentration reflects the combined influence of surface modification and sediment reworking depth, although relationships were weak and variable, scaled by biomass: shallow reworking combined with high surface modification may promote phosphate retention, whilst deeper reworking may elevate phosphate concentrations. Arrows indicate relative nutrient concentration levels. Icons illustrate the relative contribution of traits.

Collectively, these nutrient-specific patterns reveal that ecosystem functioning is spatially organised within the sediment profile. We show that distinct trait combinations cluster within surficial and deeper sediment layers to regulate discrete biogeochemical pathways. This depth-stratified structure – consistent across 17 taxonomically diverse species – demonstrates that the spatial expression of traits, rather than just their presence, is a primary determinant of species contributions to ecosystem functioning. More broadly, our findings suggest that ecosystem functioning can be defined in two dimensions, the spatial extent of environmental modification and the intensity and frequency of expression.

Importantly, this depth-stratified organisation was not captured by existing bioturbation mode typologies or taxonomic classifications. Instead, species reorganise along continuous trait axes that reflect the intensity and spatial expression of sediment reworking and ventilation processes, with ramifications for nutrient outcomes. Despite close taxonomic affinity, *Amphiura filiformis* showed markedly lower ammonium and phosphate concentrations than *A. chiajei*, driven by differences in burrow ventilation rather than taxonomy (Fig. 4, Fig. 5C and D). This aligns with previous work demonstrating that closely related benthic taxa can exhibit divergent functional contributions due to differences in trait expression and behavioural responses to environmental context, rather than taxonomic identity alone^27^. This highlights how variation in trait expression restructures functional roles even within a genus. Two polychaetes, *Nephtys caeca* and *Notomastus latericeus*, exhibited sharply contrasting bioturbation signatures and nutrient effects (Fig. 4), demonstrating that neither sediment penetration depth nor ventilation intensity alone is sufficient to predict control over specific biogeochemical pathways. Given these observations, we speculate that the same will hold true over temporal axes of determination. Understanding biodiversity-ecosystem function relationships requires disentangling intrinsic trait effects from their environmentally mediated expression, because trait-function relationships will depend on how organisms and their inherent traits interact with local environmental conditions^55,56^.

Together, these species-level relationships suggest a mechanism by which anthropogenic disturbance and climate change could alter benthic nutrient cycling if they change the abundance or expression of key traits within assemblages. Perturbations that physically alter the environment and selectively remove larger species, such as bottom trawling^39,57,58^, directly affect surface mixing and biomass, restructuring benthic communities and modifying nutrient mediation in shelf sediments^39^, whilst changes in environmental conditions, such as warming and deoxygenation, preferentially affect ventilatory behaviour^59^ and the rate and frequency of faunal processing. Hence, ecosystem change is strongest when response traits and effect traits are aligned^60^, such that the traits most sensitive to disturbance are also those exerting the strongest control over ecosystem processes. For example, larger body size has a positive effect on functioning whilst simultaneously increasing local levels of extinction risk^28^, meaning that disturbances targeting large-bodied species may disproportionately reduce key ecosystem processes. However, response and effect traits do not always align. For instance, recent work shows that high-biomass species can exert comparatively small influences on nutrient fluxes, while lower-biomass species with particular behavioural traits can dominate sediment–water exchange processes^61^. Consequently, small changes in faunal mediation caused by community reorganisation can disproportionately affect some biogeochemical components, whilst having limited effects on others^38,62^.

When the traits that respond to environmental change differ from those associated with ecosystem processes, some functions may be maintained despite shifts in species composition. In single-species incubations, nitrate no consistent relationship with measured traits (Fig. 5B).This suggests that nitrate may be buffered from in assemblage trait composition. This trait–function decoupling has been demonstrated in community level systems also. The selective alteration of feeding-guild composition by insecticides in freshwater zooplankton communities does not destabilise gross primary production because body size and omnivory, rather than mode of feeding, govern the processes that contribute most to primary production^63^. Similarly, changes in land-use can change the composition and above-ground biomass of plants, but many key ecosystem functions are associated with root morphology, leading to a weak correspondence between species turnover and functional change^64^. Sustained perturbations may shift ecosystems to alternative states^65^, particularly when the traits that make species sensitive to disturbance covary with those traits that drive ecosystem processes.

Although trait-based models predict that compensatory dynamics within functional groups may buffer species losses^21,22,38^, our results show that species assigned to the same broad functional groups can differ substantially in trait expression and nutrient responses (Fig. 4 and 5). This suggests that compensation within multispecies assemblages may depend on the intensity and spatial deployment of traits rather than group membership alone^13,19,25,37^. Species combining high surface modification with increased biomass, most notably the bivalve *Cerastoderma edule*, were associated with elevated concentrations of ammonium consistent with sediment overturning, grazing, and surface redistribution of organic material^66–68^ (Fig. 5C–D). While for some taxa, elevated ammonium concentrations may also reflect metabolic excretion rather than sediment reworking alone, our trait-based predictors capture important, though not mutually exclusive, mechanisms underlying the observed nutrient responses. The substantial within-species variability in several behavioural traits suggests that individual differences in activity or physiological condition may have contributed to unexplained variation. Only one mortality was observed and excluded, although non-recovery of some soft-bodied polychaetes introduced additional uncertainty. Upward-conveying polychaetes such as *Euclymene lombricoides* and *Notomastus latericeus* clustered within the same functional space as *C. edule* and *E*. *cordatum* despite lower biomass and only moderate ventilation, drawing together species traditionally assigned to three of the seven commonly used categorical bioturbation modes^30,68–71^. This reframes the functional role of these upward-conveying polychaetes showing that functional roles emerge from trait expression rather than taxonomic assignment or specific observed behaviours. Through sediment surface disturbance and upward particle transport (scaled by biomass), these taxa oxygenate near-surface sediments, stimulating microbially mediated remineralisation processes^72^, and strengthening bentho-pelagic coupling, elevating ammonium and phosphate production with little contribution from burrow ventilation^35,70,73,74^.

In contrast, deep-burrowing taxa exerted a fundamentally different control on nutrient pathways. Species such as *Nephtys caeca* and *Priapulus caudatus* (polychaete and priapulid worms) promote deeper sediment reworking, exposing reduced nitrogen pools to oxidised porewaters and stimulating the intermediate steps of nitrification^75^. This is reflected in increased nitrite concentrations (Fig. 5A), with nitrite subsequently transported towards the surface during burrow ventilation. At the same time, these taxa showed comparatively weak or negative effects on ammonium and phosphate concentrations, indicating retention or transformation below the sediment-water interface. These findings align with previous evidence that deep-burrow ventilation elevates redox potential, favours oxidised nitrogen forms, and suppresses ammonium and sulphide accumulation in surrounding sediments^76,77^. Mucopolysaccharide associated with the lining of their complex burrow structures may also enhance sediment nitrification rates by stimulating and fuelling nitrifying microbial groups^78^. Together, these results support a conceptual separation between taxa that regulate rapid surficial recycling and deep burrowers that act as nutrient gatekeepers, regulating both the magnitude and form of nutrient delivery to the overlying water.

Ventilation emerged primarily as a regulatory trait whose effects depended on its interaction with sediment reworking behaviour, yet ventilatory behaviour has received considerably less attention relative to mechanisms of sediment particle transport^75^. Rather than acting independently, ventilation of the burrow system increases the transport of electron acceptors into the sediment profile, leading to altered levels of nutrient generation depending on the depth and morphological characteristics of the burrow, timing, and the type of ventilation behaviour (advective vs diffusive)^24,35,62,80–82^. For the larger, deep-reworking taxa, ventilation interacts with sediment reworking depth, producing higher concentrations of nitrite. By introducing oxygen into deeper sediments, ventilation alters redox gradients and microbial activity within burrows^31,32^, regulating how and where nitrogen transformations proceed within sediments rather than uniformly stimulating nitrification^61^. This may be more widespread than appreciated, with ventilatory behaviour in the ophiuroid brittlestars *Amphiura chiajei* and *Amphiura filiformis* associated with reduced nitrite release and suppressed ammonium concentrations, acting as solute moderators rather than strong drivers. In contrast, the polychaetes *Capitellidae* sp. and *Euclymene lombricoides* showed increased rates of ventilatory behaviour with raised ammonium concentrations, indicating that, at least for some taxa, ventilatory behaviour amplifies rather than reduces net nutrient release. By integrating principal component analyses with mixed-effects models, our study helps to resolve how different configurations of bioturbation behaviour regulate distinct biogeochemical pathways, embracing inter- and intraspecific variability that is not captured in static functional group classifications.

In doing so, our analyses reveal that species contributions to ecosystem functioning are governed by both additive and interactive trait effects, the relative importance of which depends on the specific nutrient pathway. However, this trait dependency did not extend to nitrate, likely reflecting its status as a more stable and independent reservoir in marine waters^83^. These results demonstrate strong nutrient specificity, with no significant trait predictors of nitrate flux detected despite variation in species identity, biomass and bioturbation traits across the experiment. In contrast, the species-level associations of ammonium and nitrite with bioturbation traits offer a potential mechanism consistent with field observations in which shifts in community trait composition coincide with changes in dissolved inorganic nitrogen concentrations^45^. For example, in Odense Fjord, transitions toward deeper-burrowing and ventilatory taxa coincided with increased ammonium and dissolved inorganic nitrogen concentrations, consistent with the trait-nutrient relationships resolved here. Phosphate showed a different pattern, with concentrations displaying weak and variable relationships with surface modification and sediment reworking depth. Specifically, patterns suggest that phosphate associated with surface reworking may be reduced as sediments are mixed deeper, consistent with downward redistribution and sediment retention of phosphate especially as phosphate is predominantly bound within solid sediment phases^84^. These findings contrast with work elsewhere^85^ that identifies deep burrowing ventilators as the primary determinants of phosphate, which emphasises the context dependence of these pathways and the fact that different ensembles of traits become dominant under different community structures and sedimentary regimes.

Critically, these nutrient-specific differences in mechanism suggest that trait dominance can outweigh variation in trait diversity or traditional functional group categorisations^24^. Biogeochemical models often simplify faunal activity into coarse parameters such as Community Bioturbation/Bioirrigation Potential (BPc/BIPc)^86,87^, Biodiffusion coefficient (Db)^88^, faunal biomass, density^77^, or reworking depth (L)^39^ that might capture broad species-level contributions but miss specific combinations of effect-traits that cluster within defined sediment zones. Yet growing evidence shows that individual invertebrate contributions matter^19,57,89,90^. Although such proxies are useful, quantifying all invertebrates by one or two aggregate measures overlooks the diverse signals species generate, including variation attributable to intraspecific differences in relative performance, and the distinct effect traits that drive fluxes. Moreover, these effects are detectable within days of incubation, underscoring how rapidly benthic fauna can regulate bentho–pelagic coupling, and how sensitive these regulatory pathways may be under intensification of anthropogenic activity and/or climate change^91^.

Beyond nutrient fluxes, macrofaunal feeding can restructure sediment carbon quality, selectively removing labile fractions while promoting the burial of refractory material^76^. The trait combinations identified here can be resolved from standard, routinely assessed parameters. Rather than adopting static trait classifications, the approach we have adopted leverages empirically derived relationships between trait expression and nutrient pathways to parameterise predictive models. At the species level, nitrite responses were consistently associated with biomass and sediment reworking depth, suggesting that these variables may provide useful inputs for community-scale models. In contrast, ammonium concentrations emerged from an interaction between surface disturbance and ventilation, where high biomass and intense reworking promote ammonium release, moderated by increased ventilatory behaviour. For phosphate, predictions are more tentative. Given that phosphate is often strongly associated with solid sediment phases, faunal influence is likely to dependent on the capacity to physically redistribute, or expose, particle-bound phosphorus. While our observed phosphate responses were variable, the patterns we observed suggest that mobilisation is most probable when high biomass, surface modification, and greater sediment reworking depth coincide.

Unlike existing indices, such as bioturbation and bioirrigation potential^28,86^, which aggregate species contributions into fixed, additive scores, we demonstrate that these categorical proxies lose predictive power when subjected to empirical measurement (Fig. 4). Our approach, instead, utilises continuous variables of biomass and sediment reworking as direct predictors of nutrient flux. By doing so, we retain the inherent interaction structure between traits, preserving the critical mechanistic information that categorical indices omit. For example, identifying taxa that combine high surface modification with substantial biomass, or deep-burrowing, ventilating taxa, may allow managers to move beyond simple species lists toward safeguarding the functional drivers of benthic biogeochemistry. Protecting ecosystem functioning may therefore require consideration of the expression and balance of key trait combinations alongside taxonomic diversity. Our species-level findings demonstrate that nutrient responses are contingent on specific trait combinations, suggesting that perturbations altering their spatial distribution and intensity could reorganise biogeochemical pathways in ways consistent with field observations..

Our findings matter within the current discourse about harnessing ecosystems to help with rapid adaptation to, and mitigation of, the effects of intensifying anthropogenic activity ^91^ and climate change^92^. Examples of scalable good practices that capitalise on the linkage between biodiversity and functioning exist^93^, but challenges remain in determining how, when and where the benefits derived from socio-ecological systems arise^94^. From a functional standpoint, key trait– environment dynamics can be more important than biodiversity^95^ in maintaining ecosystem processes and functions, with profound consequences for nutrient cycling, carbon storage and ecosystem resilience^46,96^. Our results identify species-level trait mechanisms associated with variation in benthic nutrient concentrations and therefore support the broader argument that conservation assessments should consider functional traits and processes alongside species inventories. As nations commit to protecting 30% of land and sea by 2030^97^ under the Kunming– Montreal Global Biodiversity Framework, conservation must extend beyond these inventories to the functional processes that confer ocean resilience in an era of accelerating disturbance^91^.

## Methods

### Invertebrate fauna and sediment collection

Sediment and invertebrate fauna were collected across 11 stations off the Isle of Cumbrae, Scotland using a 0.1m^2^ Day grab deployed from the *r.v. Actinia* in April 2023 (Supplementary Table S1). Sediment was sieved (500 μm mesh) in a seawater bath to remove macrofauna and debris, allowed to settle for 24 h to retain the fine fraction (less than 63 μm) and stirred to homogenize the distribution of particles (mean ± SD, n = 3: median particle size, d50, 86.57 ± 2.3 µm; range, d10-d90, 8.12 ± 1.7 - 205.33 ± 20.5 µm; organic matter content, 4.43 ± 0.11 %). We collected 17 invertebrate species across 5 major taxonomic groups (<u>Bivalvia</u>, *Abra nitida* (glossy furrow shell), *Cerastoderma edule* (common cockle), *Mysia undata* (wavy venus), *Phaxas pellucidus* (transparent razor shell); <u>Gastropoda</u>, *Turritellinella tricarinata* (auger shell); <u>Polychaeta</u>, *Capitellidae sp.* (gallery worm), *Euclymene lombricoides* (bamboo worm), *Glycera alba* (bloodworm), *Lagis koreni* (trumpet worm), *Nephtys caeca* (catworm), *Notomastus latericeus* (bristleworm), *Terebellides stroemii* (terebellid worm); <u>Echinodermata</u>, *Amphiura chiajei* (brittlestar), *Amphiura filiformis* (brittlestar), *Echinocardium cordatum* (heart urchin), *Paraleptopentacta elongata* (sea cucumber); and <u>Priapulida</u>, *Priapulus caudatus* (cactus worm)) by hand from sieve (2mm mesh) returns. Individuals were held in glass aquaria for 48 h in aerated, in natural seawater only at 12 °C under a 16:8 h light:dark cycle and were not fed prior to the experiment starting.

### Experimental set-up

We introduced a single individual of each invertebrate species to replicate (n = 5) transparent acrylic aquaria with dimensions (L × W × H: small, 2.2 × 2.2 × 25 cm; large, 6 × 6 × 25 cm) appropriate for each species body size and activity patterns (small aquaria: Bivalvia, *A. nitida*, *M. undata*, *P. pellucidus*; Polychaeta, *Capitellidae sp.*, *E. lombricoides*, *L. koreni*, *N. latericeus*; Echinoderms, *A. chiajei*, *A. filiformis*; large aquaria: Bivalvia, *C. edule*; Gastropoda, *T. tricarinata*; Polychaeta, *G. alba*, *N. caeca*, *T. stroemii*; Echinodermata, *E. cordatum*, *P. elongata*; Priapulida, *P. caudatus*). To quantify the contribution of the microbial and meiofaunal community on our response variables, we also assembled replicate (n = 5) small and large aquaria containing no macrofauna. Each aquarium contained sieved sediment to a depth of ∼13 cm and seawater (salinity 33ppt) to ∼10.5 cm (equivalent to: small aquaria, ∼57 ml; large aquaria, 410 ml) above the sediment-water interface. Aquaria were placed randomly across three temperature-controlled water baths (12 ± 0.5 °C, Teco TK 2000 chiller/heater) under a 16:8 blue light-dark cycle (Aquabar T-series blue LED, 450 nm). All aquaria were continually aerated and the experiment ran for 9 days. After the first 24 hours 75% of the overlying water was replaced to remove excess nutrients associated with assembly..Target replication was n = 5 per species for nutrient measurements. As some individuals were not recovered for post-incubation biomass measurement, replication in the trait-nutrient models was reduced for *Capitellidae* sp., *E. lombricoides* and *N. latericeus* (n = 2), *T. stroemii* (n = 3), and *N. caeca* and *C. edule* (n = 4); all other species were represented by n = 5. Analyses used all available replicates, with species identity modelled as a random effect. Individual biomass (g, wet weight) was measured post-incubation after gently blotting surface water; bivalves and gastropods were weighed with shells; echinoderms and worms were weighed whole. Failure to recover an individual at the end of the experiment was treated as missing biomass rather than evidence of mortality, because all unrecovered individuals were soft-bodied polychaetes that may fragment, remain concealed within sediment or be lost during recovery and sieving. The single observed mortality of *C. edule* was excluded from all analyses.

### Faunally-mediated sediment particle reworking

Faunally mediated sediment particle reworking was estimated from the vertical redistribution of an optically distinct particle tracer (fluorescent green sand: median particle size, 317 µm; range d10-d90, 219-452 µm; Glass Pebbles Ltd., UK) using fluorescent-Sediment Profile Imaging (f-SPI^98^). Particle tracers were evenly spread across the sediment surface at a concentration of ∼0.65 g cm^−2^ (small aquaria, 4 g aquarium^−1^; large aquaria, 20g aquarium^−1^) after 12 hours of incubation allowing the organisms to burrow and, after 9 days, images of the four sides of each aquarium were taken using a digital SLR camera (Canon 400D, 15 s exposure, f5.6, ISO400, effective resolution = 81 µm pixel^−1^) housed within a UV illuminated box (Schiffers et al.^99^). Images were stitched together (JPEG compression, Supplementary Figs. S5-23) for each aquarium, and particle tracer depth profiles were generated (Supplementary Figs. S26 and S27) using a customized script in ImageJ (v. 1.47 s), a java-based public domain program developed at the US National Institutes of Health (https://imagej.net/ij/index.html). From these data, we calculated the mean (^f-SPI^L_mean_, typical short-term depth of mixing) and maximum (^f-SPI^L_max_, maximum extent of mixing over the long-term) mixed depth of particle redistribution and the maximum vertical deviation of the sediment-water interface (upper – lower limit surface boundary roughness, ^f-SPI^SBR, an indication of surficial activity^25^. In addition, using plan-view images of the sediment surface (sediment surface imagery, ^SSI^S_mod_; Supplementary Figs. S24, and S25), we estimate the extent of surface modification of particles from depth to the sediment surface (^SSI^S_mod_, cm^2^) due to conveyor belt, ploughing or other accelerated transport mechanisms.

### Burrow ventilation

We estimated the ventilatory behaviour of each individual from changes in the concentration (∼10 mM: small aquaria, 0.049 g Br aquaria^−1^; large aquaria, 0.370 g NaBr aquaria^−1^) of the inert tracer sodium bromide Δ[Br⁻]; calculated as the difference between bromide concentrations measured at 8 h and 0 h ([Br⁻]₈ₕ - [Br⁻]₀ₕ), expressed in mg L⁻¹. Sodium bromide was added to each aquarium as a prepared 20ml aqueous solution and the overlying water was gently mixed before a 12 ml sample was collected as the 0 h measurement. A second 12 ml sample was collected after 8 h. Samples were filtered through 0.7 µm GF/F filters, and bromide concentrations were determined using a Tecator flow-injection auto-analyser (FIA Star 5010 series). Negative values indicated greater removal of bromide from the overlying water through faunal ventilation.

### Water column nutrient concentrations

Accumulated water column nutrient concentrations (Ammonia, NH_4_-N; Nitrite, NO_2_-N; Nitrate, NO_3_-N; and Phosphate, PO_4_-P; µmol L^−1^) were quantified after 8 days from standardized water samples (5 cm water depth, 0.45 µm filtered and frozen prior to analysis) following standard colorimetric methods (Grasshoff et al.^100^) using a Lachat Quikchem 8500 flow-injection auto-analyser. These measurements therefore represent final water-column concentrations integrating the net outcome of faunal, microbial and abiotic processes during the incubation.

### Log-Normal Response Ratios

Because aquarium dimensions, overlying-water volume and headspace differed between the two aquarium sizes, direct comparison of raw response variables was not appropriate. We therefore expressed each faunal treatment relative to the no-macrofauna controls of the corresponding aquarium size using the log normal response ratio (LNRR): LNRR = ln(mean treatment/mean control). Positive LNRR values indicate an increase relative to the corresponding control, whereas negative values indicate a decrease. This standardisation accounts for size-specific differences in baseline nutrient concentrations and process measurements. As our experimental design required that organisms were housed in two sizes of aquaria, we cannot exclude the effect of aquarium geometry on species behaviour, although previous work has shown these effects are minimal^101^. The LNRR transformation does not exclude potential effects of aquarium dimensions on faunal behaviour, sediment reworking, ventilation or solute exchange. Biomass was not standardised using LNRR and was included as a separate covariate in the statistical analyses.

For nutrients, LNRR compared the final concentration in each faunal treatment with the mean final concentration in no-macrofauna controls of the corresponding aquarium size. Positive values therefore indicate higher endpoint concentrations than the corresponding controls, whereas negative values indicate lower endpoint concentrations.

### Dimensionality reduction

As the mechanistic basis of faunal contributions to ecosystem functioning often correlate with one another, we used principal component analysis (PCA) to reduce the dimensionality of our set of explanatory variables (^f-SPI^SBR, ^SSI^S_mod_, ^f-SPI^L_mean_, ^f-SPI^L_max_, [Br−]) using the *stats* (v 4.2.2) function in R^102^. Correlated variables were identified from visual inspection of the PCA biplot (Supplementary Fig. S1) and confirmed using Pearson correlations (when r >50%, Supplementary Fig. S2). Decisions on which explanatory variable to remove from each correlated pair were based on relevance to nutrient cycling. Multicollinearity was investigated using generalized variance inflation factors using the VIF function in the *car* package in R. All VIFs were less than 4.

### Intraspecific variability in species contributions

To assess the consistency of faunal contributions to sediment particle reworking, we assessed the intraspecific trait variability (ITV, De Bello et al.^103^, Equation 1) for all of our explanatory variables. For each explanatory variable, we fitted a one-way ANOVA with species identity as the explanatory factor. Intraspecific trait variability was calculated as the percentage of the total variation attributable to within-species variation:

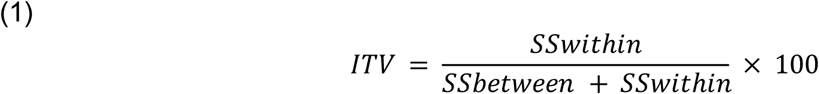

where *SS_between_* is the sum of squares associated with species identity and *SS_within_* is the residual sum of squares. Higher ITV values indicate greater within-species variability relative to between-species differences, whereas lower values indicate stronger species-level structuring of trait expression.

### Relationship between trait expression and nutrient release

We used linear mixed effects models - to analyse trait–nutrient relationships (a robust approach for complex ecological datasets^104^). For each nutrient response (NH₄⁺-N, NO₂⁻-N, NO₃⁻-N, PO₄³⁻-P; all expressed as LNRR), we fitted the same global model: *response* ∼ biomass (g) * ^SSI^S_mod_ (LNRR)* ^f-SPI^L_max_ (LNRR) * Δ[Br^−^] (LNRR) + (1 | species_name) (lme4/lmerTest). Biomass was not LNRR-standardised and entered as a separate covariate. As our focus was to establish the mechanistic basis of faunal contributions to nutrient release, and not to determine differences among individual species, a random intercept for species identity (1|species_name) was included. Models were fitted by maximum likelihood (REML = FALSE) to enable comparison during selection. Prior to modelling, we reduced redundancy among traits via PCA and Pearson correlations; among retained predictors, multicollinearity was checked with *car::vif* (all VIF < 3). We applied backward elimination with *lmerTest::step*: the random structure was evaluated using likelihood-ratio tests, and fixed terms were removed sequentially using Satterthwaite F-tests, with AIC/BIC inspected to confirm parsimony. Final, nutrient-specific models comprise the subset of main effects and interactions retained by this procedure. To assess whether support for the ammonium surface modification × ventilation interaction was a result of the backward elimination, we compared the retained interaction model with an otherwise identical additive model in which the interaction was removed, using a likelihood-ratio test and AIC. Model fit was summarised with marginal and conditional R² (*performance::r2*), and species-level variance contributions were expressed as an inter-class correlation coefficients calculated using *VarCorr*. All analyses were conducted in R (v4.2.2) using *lme4*, *lmerTest*, *car*, and *performance*.

Model assumptions were assessed by visual inspection of residuals versus the fitted values to evaluate homoscedasticity, and quantile-quantile plots to assess normality. Residuals were evenly distributed around zero, and no substantial deviations from normality were observed.

To quantify the proportion of total variance attributable to species identity, we calculated the intraclass correlation coefficient (ICC) for each final mixed-effects model from the estimated variance components. The ICC was computed as the ratio of the species-level random-intercept variance to the total variance (species-level variance + residual variance). This provides a measure of the degree to which unexplained variation in nutrient concentrations is structured at the species level after accounting for fixed effects. ICC was preferred over dispersion metrics (e.g. standard deviation) because it explicitly partitions variance within the hierarchical mixed-effects model and allows direct comparison of species-level structuring across models.

We visualised marginal effects from the final mixed-effects models using ggeffects::ggpredict and ggplot2 (packages: *ggeffects*, *ggplot2*, *dplyr*, *patchwork*). For each panel, we varied a focal predictor on the x-axis and held moderators at representative levels (mean and ±1 SD); lines show fitted values and ribbons 95% CIs, with interactions shown via facets or colour. The focal predictor for each nutrient was the most influential fixed term in the final model based on likelihood-ratio *χ*² tests.

To assess sensitivity to low species-level replication, the final ammonium, nitrite and phosphate models were refitted after excluding the three species represented by two complete observations (*Capitellidae* sp., *E. lombricoides* and *N. latericeus*). Fixed-effect estimates from these models were compared with those obtained from the full dataset. Nitrate was not included because its final model retained no trait predictors (Sensitivity analyses can be found in Supplementary Table S2).

### Species contributions to models (Proportion of variance)

For each final model we extracted the random-intercept conditional modes for species using *ranef*, squared them to obtain variance-scale contribution scores, and normalised within model so the species-level shares summed to 1. These proportions indicate which species account for the largest share of the between-species random-effect variance after controlling for fixed effects.

### Species specific residuals within the models

For each final mixed-effects model, we computed response residuals from the fitted object (*lme4*). We paired residuals with the corresponding species factor and, for each species, summarised the distribution and median residual. To visualise potential systematic under- or over-prediction after accounting for fixed and random effects, we plotted species-wise residual distributions as boxplots with overlaid points. Species were ordered by their median residual to aid comparison. This procedure was applied identically to all nutrients.

## Supporting information

Supplementary Inofrmation

## Ethics Approval

Ethical approval was obtained from the Biosciences Research Ethics Committee (Application ID 1825511) at the University of Exeter to undertake this work.

## Data availability

The research data supporting this publication are openly available from Harvard Dataverse at: https://doi.org/10.7910/DVN/WT89GZ.

## Acknowledgements

This work was funded by the Convex Seascape Survey (https://convexseascapesurvey.com/). The authors would like to thank all the members of the survey for their contributions to the inception and interpretation of the data herein. We thank the captain and crew of the R/V Actinia and the team at FSC Millport for their support in logistics and planning. Thanks also to Carter Melnick, Cassia Wilson and Naomi Hart for their assistance in laboratory construction, sample collection, and sampling, and to Naomi Hart for her stunning artwork that has helped communicate our findings to a wide audience.

## Author Contributions

A.P.: Conceptualisation, Methodology, Validation, Formal analysis, Investigation, Data curation, Writing – original draft, Writing – review & editing, Visualisation.

M.S.: Conceptualisation, Methodology, Validation, Formal analysis, Writing – original draft,

Writing – review & editing, Visualisation.

J.A.G.: Conceptualisation, Methodology, Validation, Formal analysis, Writing – original draft, Writing – review & editing, Visualisation.

V.K.: Writing – review & editing, Validation.

M.F.: Investigation, Data curation, Writing – review & editing. TW: Investigation, Data curation, Writing – review & editing. C.R.: Writing – review & editing, Funding Acquisition.

C.L.: Conceptualisation, Methodology, Validation, Data curation, Writing – original draft, Writing

– review & editing, Visualisation, Funding Acquisition.

