## Supplementary Inofrmation for "Sediment depth-dependent trait expression controls species contributions to nutrient cycling"

**Supplementary Information**

**Supplementary Table S1 - Sample Sites**

| *Station*  *Number* | *Depth (m)* | *Lat DDM* | *Lon DDM* | *Lat DD* | *Lon DD* |
| --- | --- | --- | --- | --- | --- |
| *1* | 42 | 55° 45.637 N | 004° 52.624 W | 55.760617 | -4.877067 |
| *2* | 30 | 55° 45.890 N | 004° 52.320 W | 55.764833 | -4.872 |
| *3* | 28 | 55° 45.200 N | 004° 52.560 W | 55.753333 | -4.876 |
| *4* | 45 | 55° 45.571 N | 004° 53.524 W | 55.759517 | -4.892067 |
| *5* | 43 | 55° 44.448 N | 004° 54.981 W | 55.7408 | -4.91635 |
| *6* | 57 | 55° 43.992 N | 004° 55.593 W | 55.7332 | -4.92655 |
| *7* | 57 | 55° 43.252 N | 004° 55.246 W | 55.720867 | -4.920767 |
| *8* | 43 | 55° 42.404 N | 004° 55.598 W | 55.706733 | -4.926633 |
| *9* | 101 | 55° 42.676 N | 004° 58.502 W | 55.711267 | -4.975033 |
| *10* | 114 | 55° 44.250 N | 004° 59.000 W | 55.7375 | -4.983333 |
| *11* | 32 | 55° 44.333 N | 004 56.400 W | 55.738883 | -4.94 |

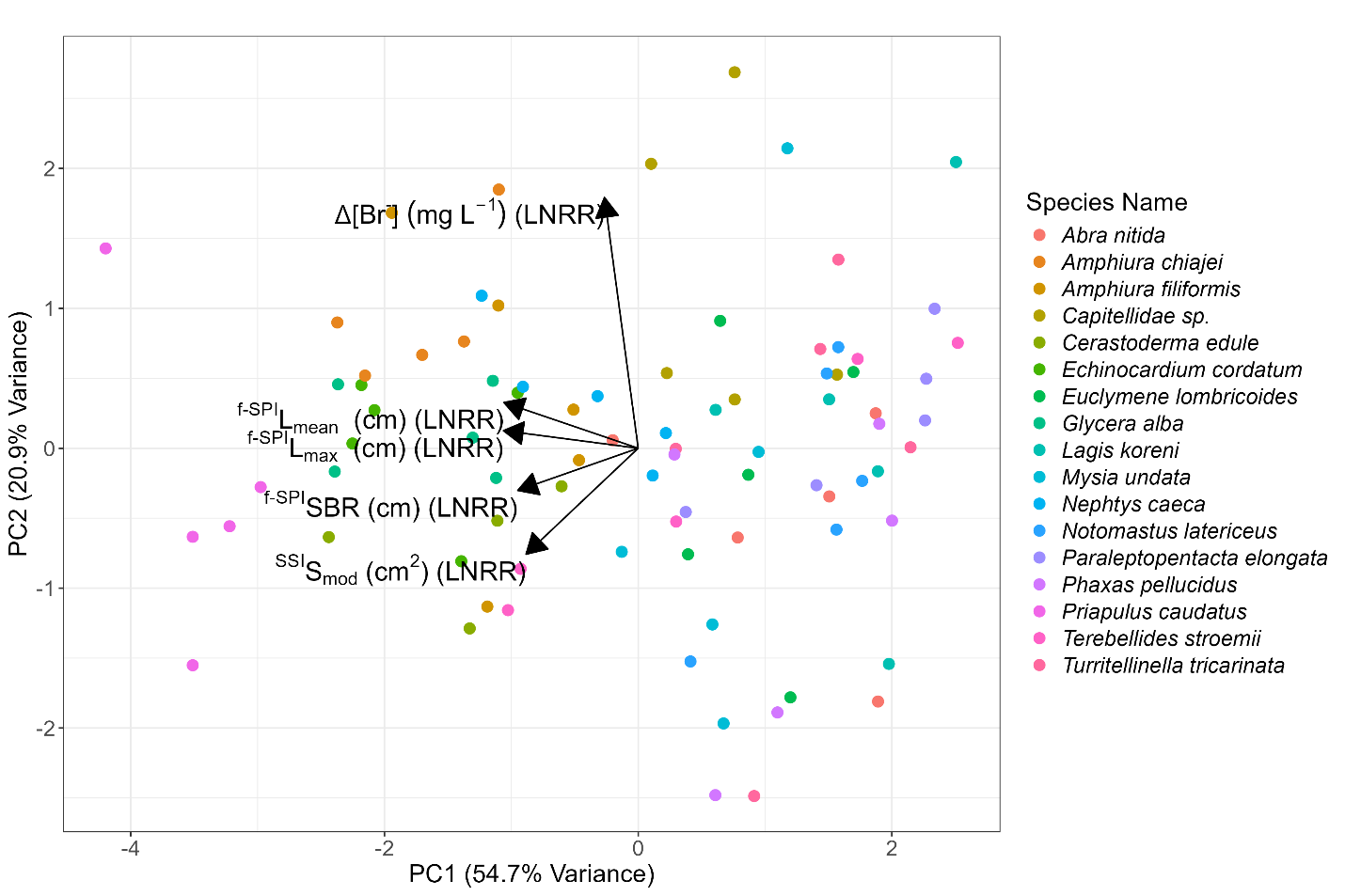

**Figure S1. Principal Component Analysis (PCA) of log response ratios (LNRR) of bioturbation traits across benthic invertebrate species.** Arrows indicate PCA loadings of explanatory variables. Arrow length reflects the strength of correlation between variables and the principal components (longer arrows = stronger correlation). Arrow direction relative to the axes shows whether variables positively or negatively influence the principal components. Variables with arrows pointing in similar directions are positively correlated, those in opposite directions are negatively correlated, and perpendicular arrows indicate no correlation.

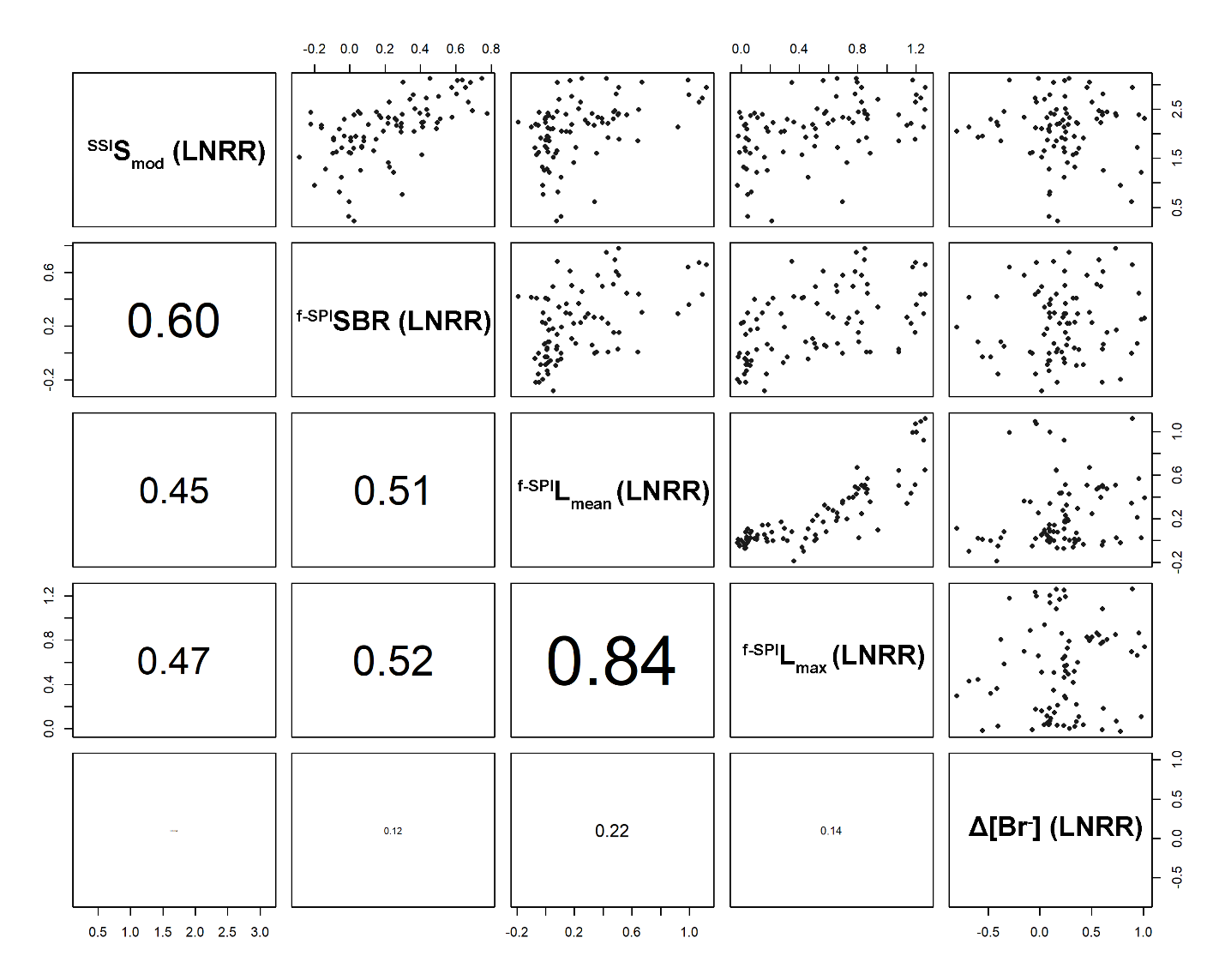

**Figure S2 Cleveland dotplot matrix of pairwise trait correlations.** Cleveland dotplots (upper panels) and Pearson’s correlation coefficients (lower panels) show relationships among log response ratios (LNRR) of surface modification (^SSI^S_mod_), sediment boundary roughness (^f-SPI^SBR), mean sediment reworking depth (^f-SPI^L_mean_), maximum sediment reworking depth (^f-SPI^L_max_), and ventilatory activity (Δ[Br⁻]). Correlation coefficients are scaled by font size, with stronger correlations displayed more prominently.

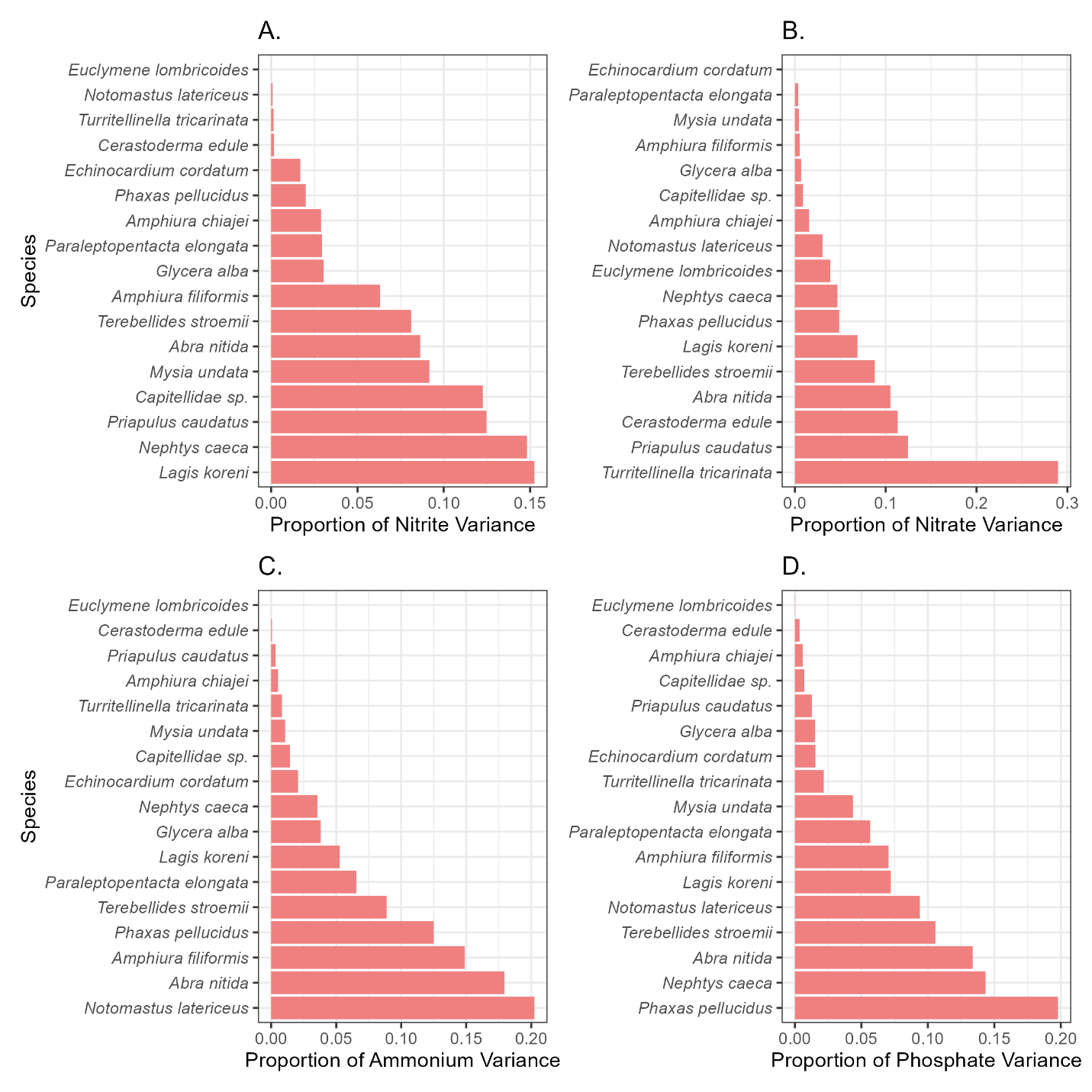

**Figure S3. Species-specific contributions to model variance for each nutrient.** Bars show the proportion of variance explained by each species in models of (A) nitrite, (B) nitrate, (C) ammonium, and (D) phosphate. Higher values indicate greater variability or uncertainty in the mean estimate (larger coefficient of variation, CV), whereas lower values indicate more precise estimates. This highlights both inter- and intra-specific differences in the strength and consistency of species’ contributions across nutrients.

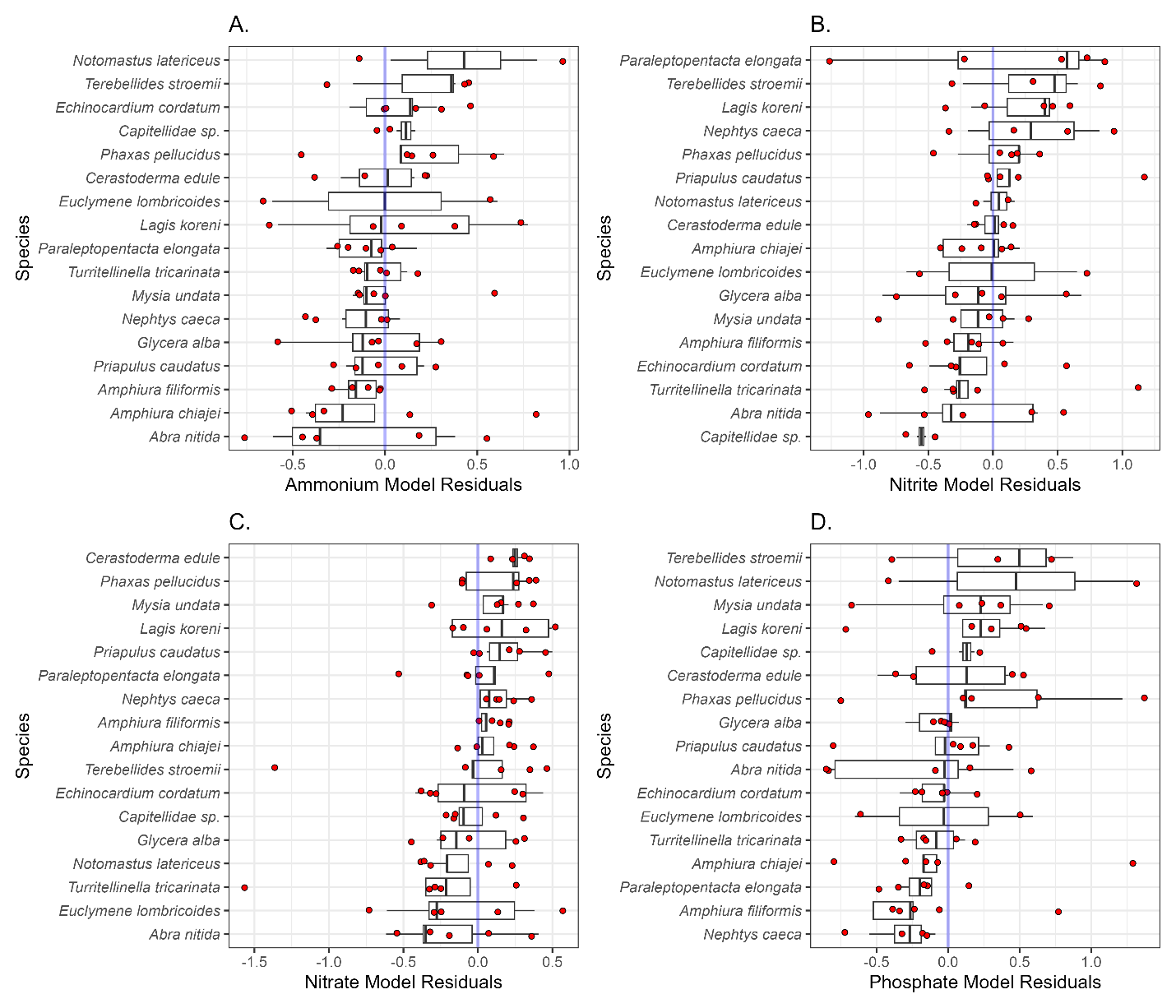

**Figure S4. Species-specific residuals from nutrient flux models.** Boxplots show the distribution of residuals for each species in models of (A) ammonium, (B) nitrite, (C) nitrate, and (D) phosphate, with individual observations plotted as red points. The blue vertical line indicates zero (perfect model fit). Residual spread around zero reflects the extent to which model predictions captured species-level variation, with wider distributions indicating greater unexplained variability and consistent deviations from zero highlighting species-specific bias in model fit.

| **Nutrient** | **Predictor** | **Full model** | **Excluding n=2 species** |
| --- | --- | --- | --- |
| Ammonium | *Biomass* | *0.045 (0.017),*  *p 0.014* | *0.050 (0.016),*  *p 0.005* |
| Ammonium | *Surface modification* | *0.403 (0.107),*  *p <0.001* | *0.406 (0.101),*  *p <0.001* |
| Ammonium | *Ventilatory activity* | *1.164 (0.473),*  *p 0.016* | *0.973 (0.516),*  *p 0.064* |
| Ammonium | *Surface modification x ventilatory activity* | *-0.583 (0.223),*  *p 0.011* | *-0.527 (0.240),*  *p 0.032* |
| Nitrite | *Biomass* | *0.047 (0.017),*  *p 0.012* | *0.044 (0.017),*  *p 0.017* |
| Nitrite | *Maximum sediment reworking depth* | *0.588 (0.174),*  *p 0.002* | *0.591 (0.178),*  *p 0.003* |
| Phosphate | *Biomass* | *-0.713 (0.496),*  *p 0.156* | *-0.358 (0.492),*  *p 0.470* |
| Phosphate | *Surface modification* | *0.355 (0.212),*  *p 0.099* | *0.348 (0.198),*  *p 0.084* |
| Phosphate | *Maximum sediment reworking depth* | *0.663 (0.914),*  *p 0.471* | *0.942 (1.025),*  *p 0.363* |
| Phosphate | *Biomass * surface modification* | *0.252 (0.184),*  *p 0.176* | *0.133 (0.182),*  *p 0.468* |
| Phosphate | *Biomass * maximum reworking depth* | *1.289 (0.777),*  *p 0.102* | *0.754 (0.776),*  *p 0.335* |
| Phosphate | *Surface modification * maximum reworking depth* | *-0.648 (0.410),*  *p 0.119* | *-0.632 (0.447),*  *p 0.163* |
| Phosphate | *Biomass * surface modification * maximum reworking depth* | *-0.422 (0.277),*  *p 0.134* | *-0.243 (0.276),*  *p 0.381* |

**Supplementary Table S2. Sensitivity of trait–nutrient relationships to low species-level replication.** Fixed-effect estimates (±SE), and associated p-values are shown for the final models fitted to the full dataset and after excluding the three species represented by two complete observations (*Capitellidae sp., Euclymene lombricoides and Notomastus latericeus*) in each trait-nutrient model. The same fixed- and random-effects structures were retained, and sensitivity models contained 66 observations across 14 species. Nitrate is not shown because its final model contained no trait predictors.

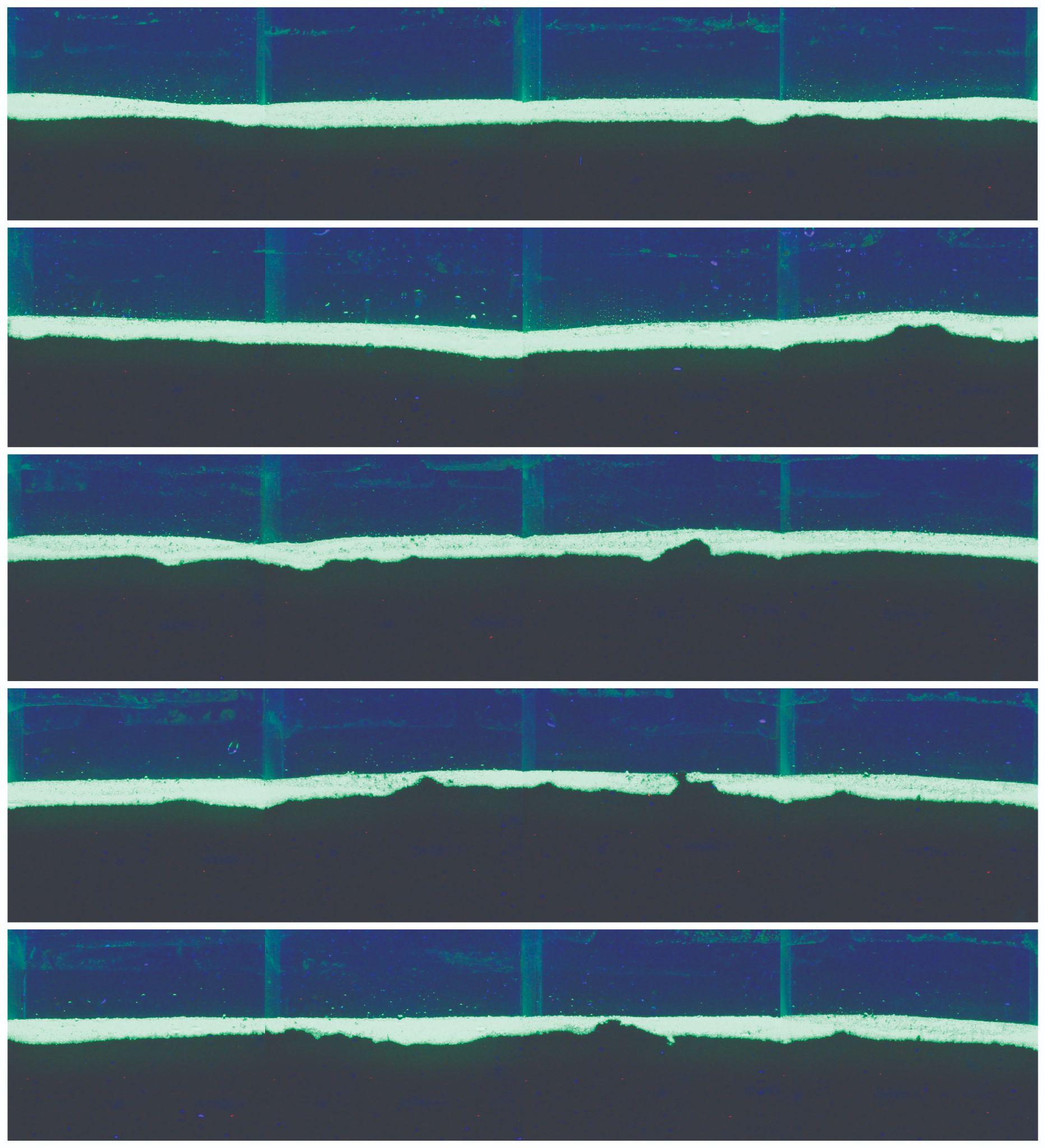

**Figure S5. f-SPI Images for 6 x 6 cm control cores with no macrofauna in.** The five images show the five replicate cores.

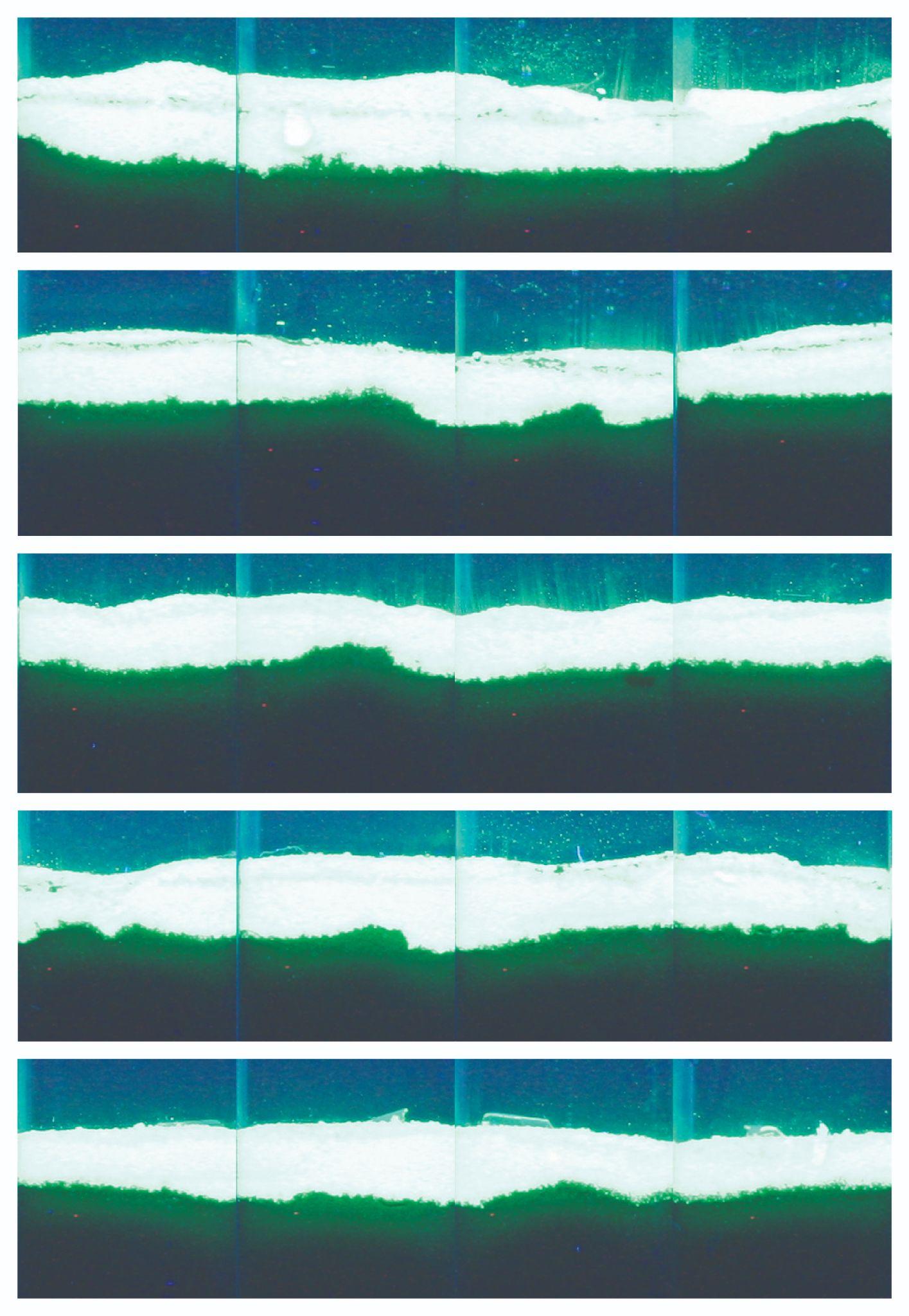

**Figure S6. f-SPI Images for 2.2 x 2.2 cm control cores with no macrofauna in.** The five images show the five replicate cores.

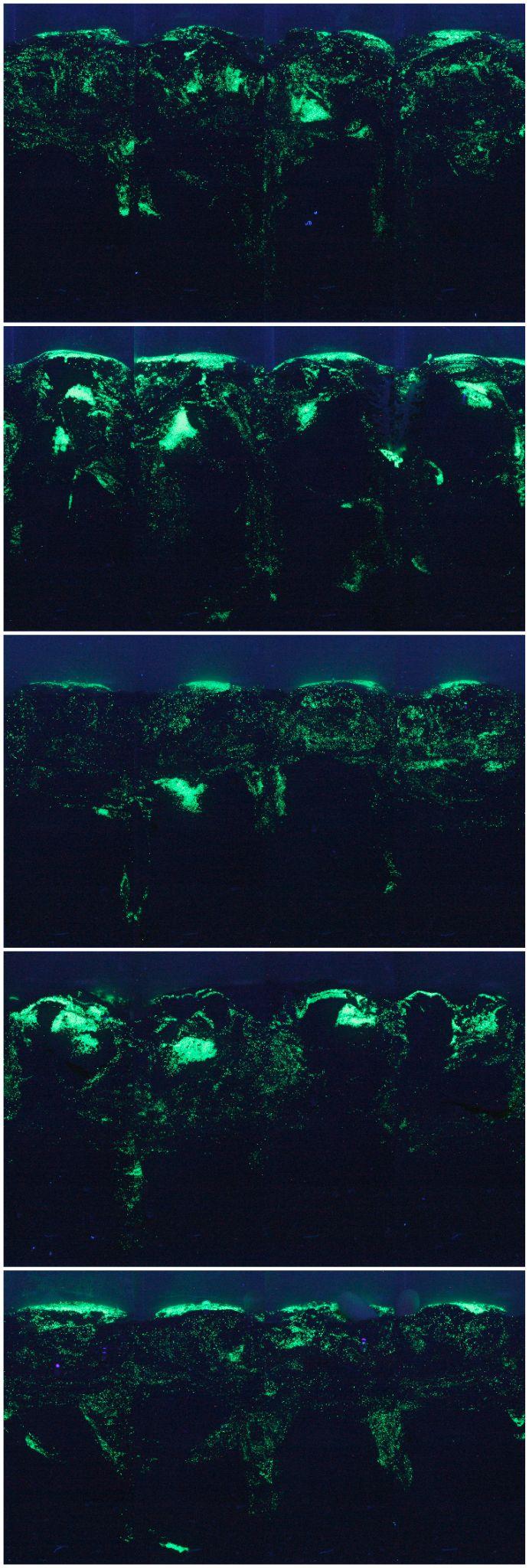

**Figure S7. f-SPI Images for *Priapulus caudatus* in 6 x 6 cm cores.** The five images show the five replicate cores.

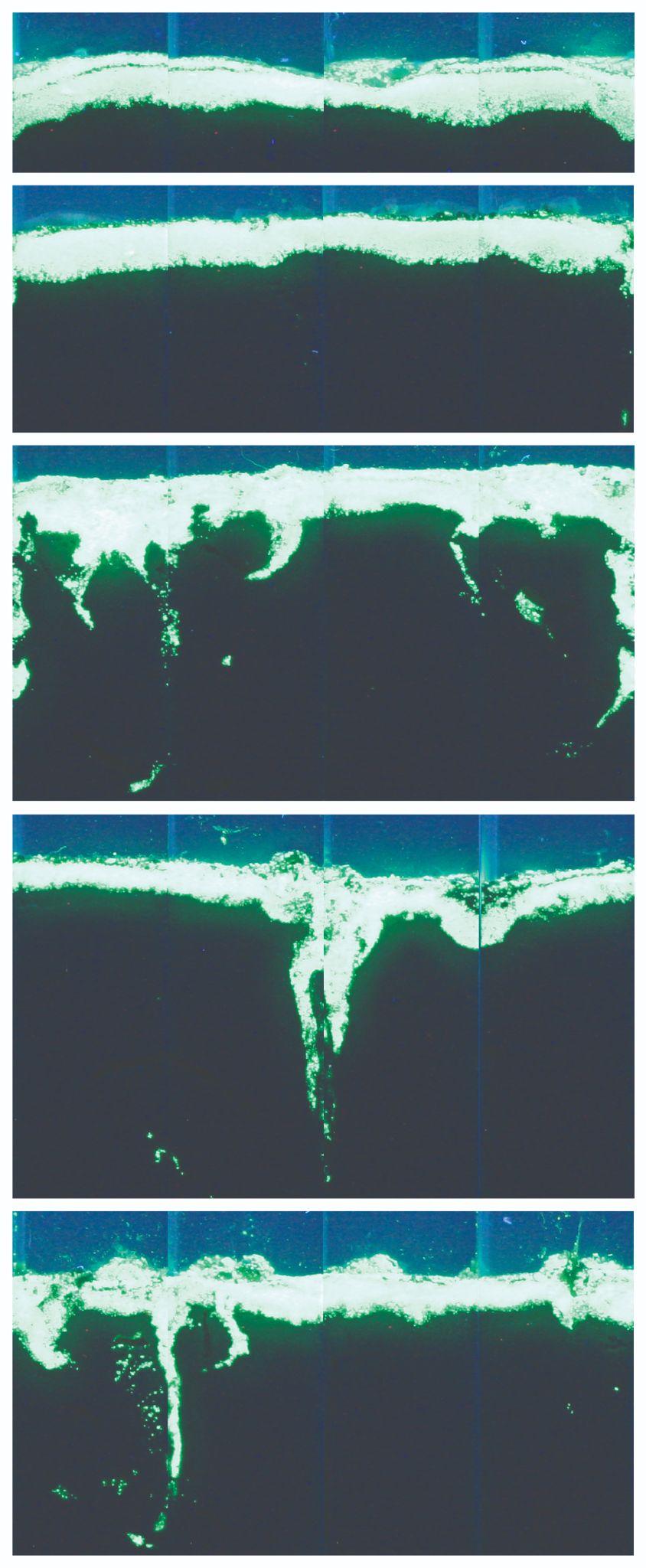

**Figure S8. f-SPI Images for *Capitellidae sp.* in 2.2 x 2.2 cm cores.** The five images show the five replicate cores.

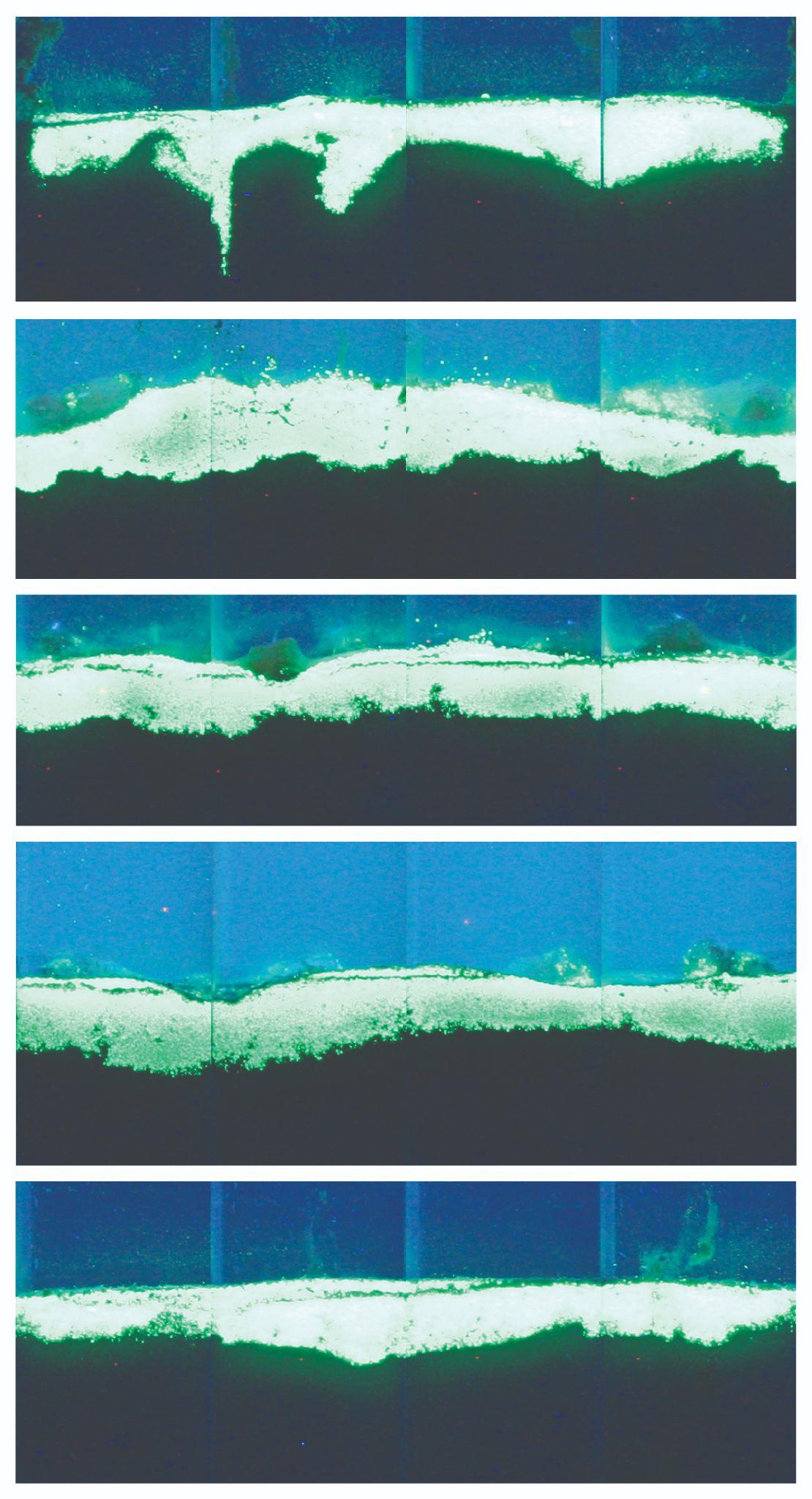

**Figure S9. f-SPI Images for *Euclymene lombricoides* in 2.2 x 2.2 cm cores.** The five images show the five replicate cores.

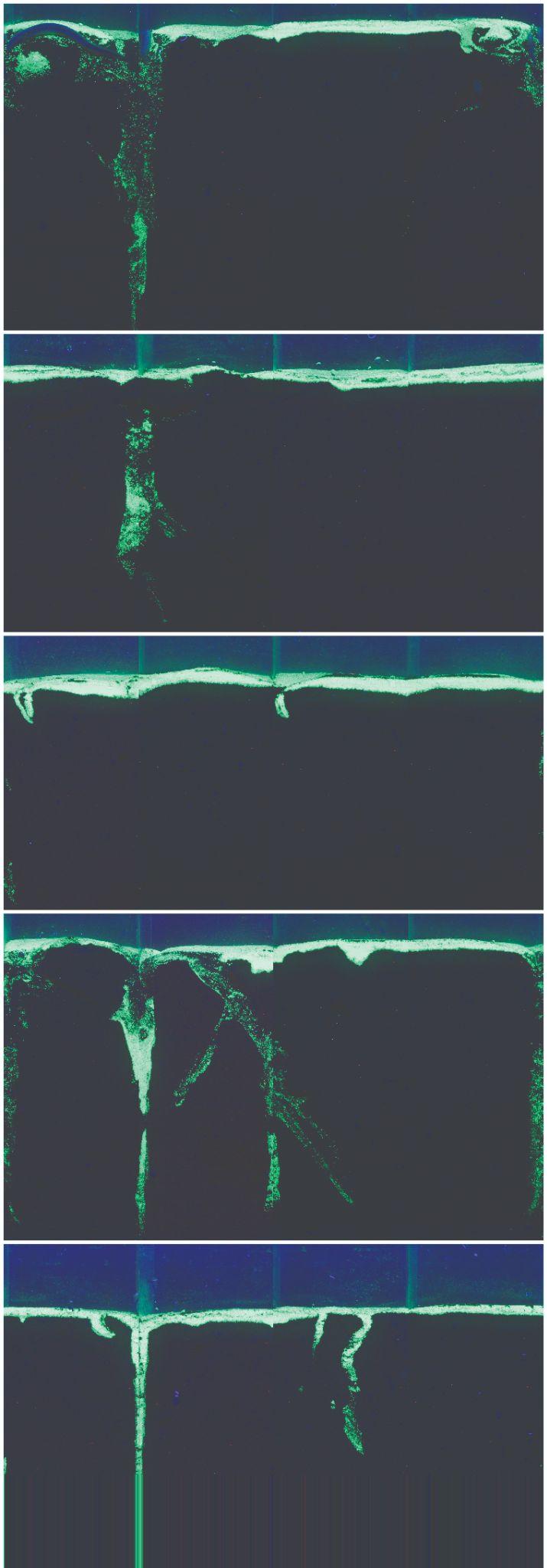

**Figure S10. f-SPI Images for *Glycera alba* in 6 x 6 cm cores.** The five images show the five replicate cores.

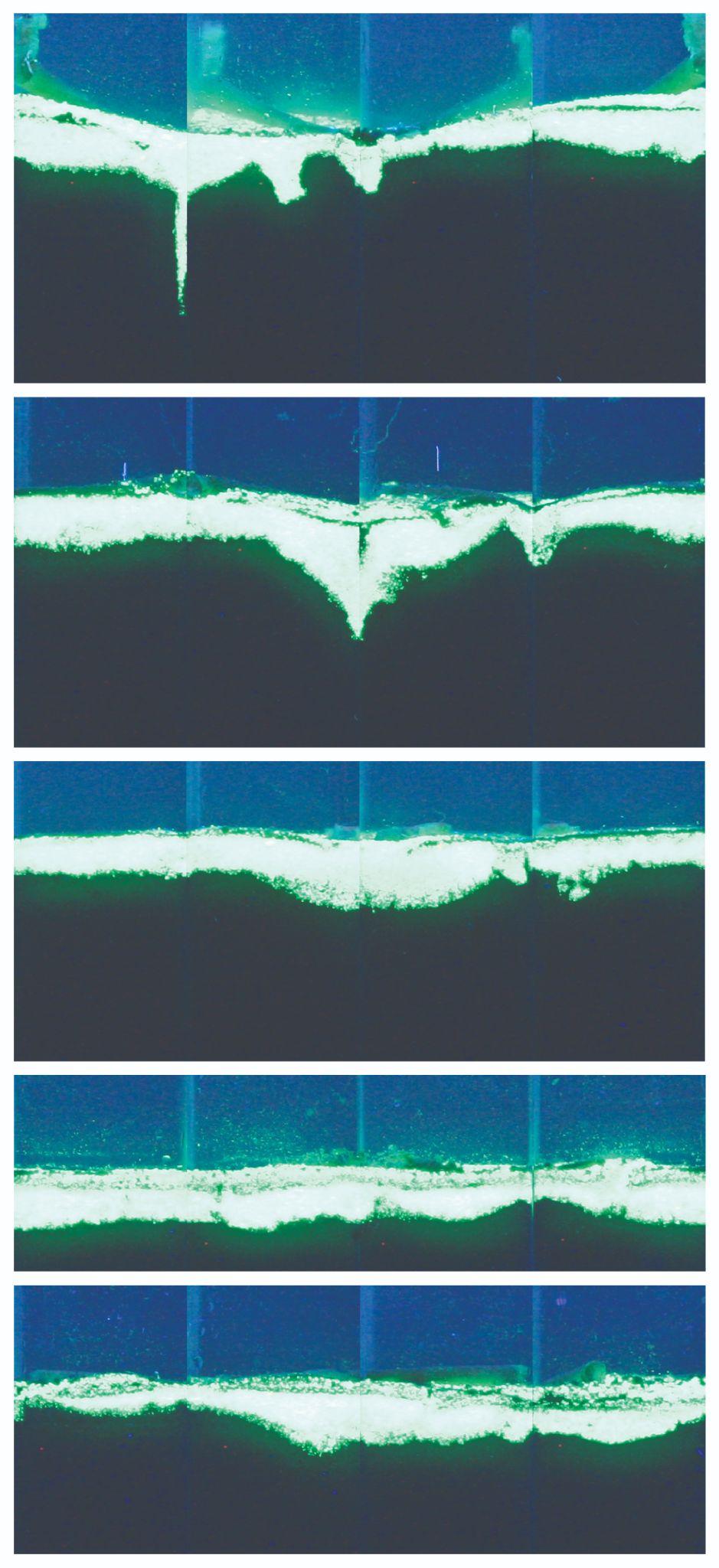

**Figure S11. f-SPI Images for *Lagis koreni* in 2.2 x 2.2 cm cores.** The five images show the five replicate cores.

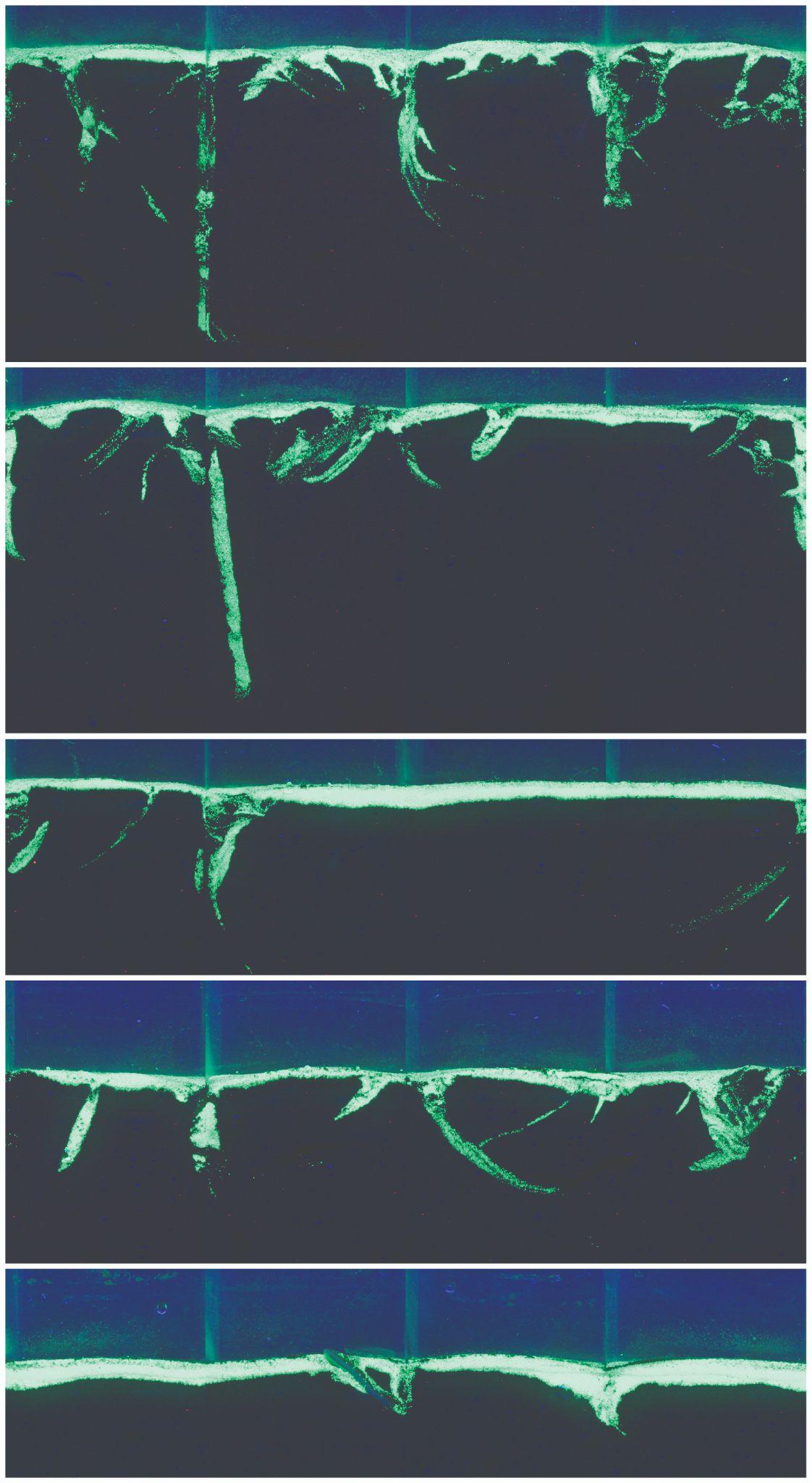

**Figure S12. f-SPI Images for *Nephtys caeca* in 6 x 6 cm cores.** The five images show the five replicate cores.

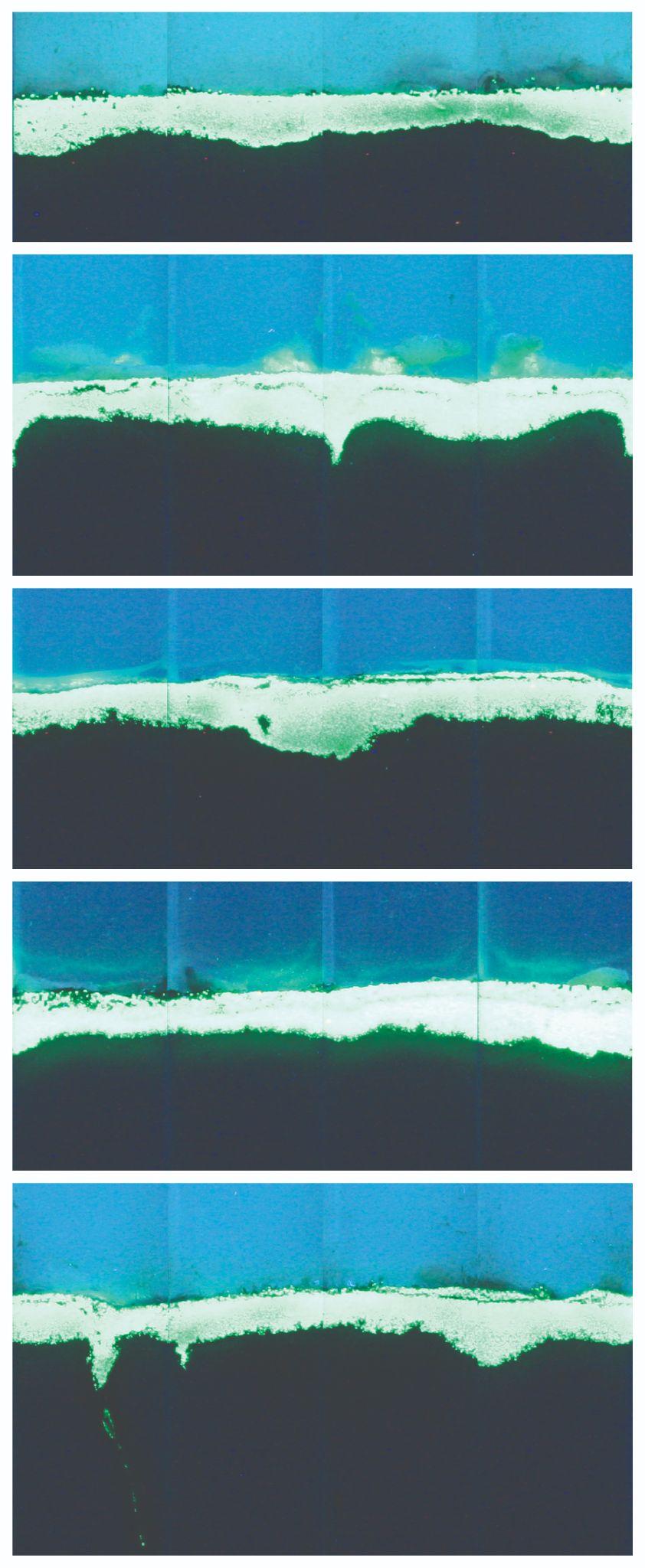

**Figure S13. f-SPI Images for *Notomastus latericeus* in 2.2 x 2.2 cm cores.** The five images show the five replicate cores.

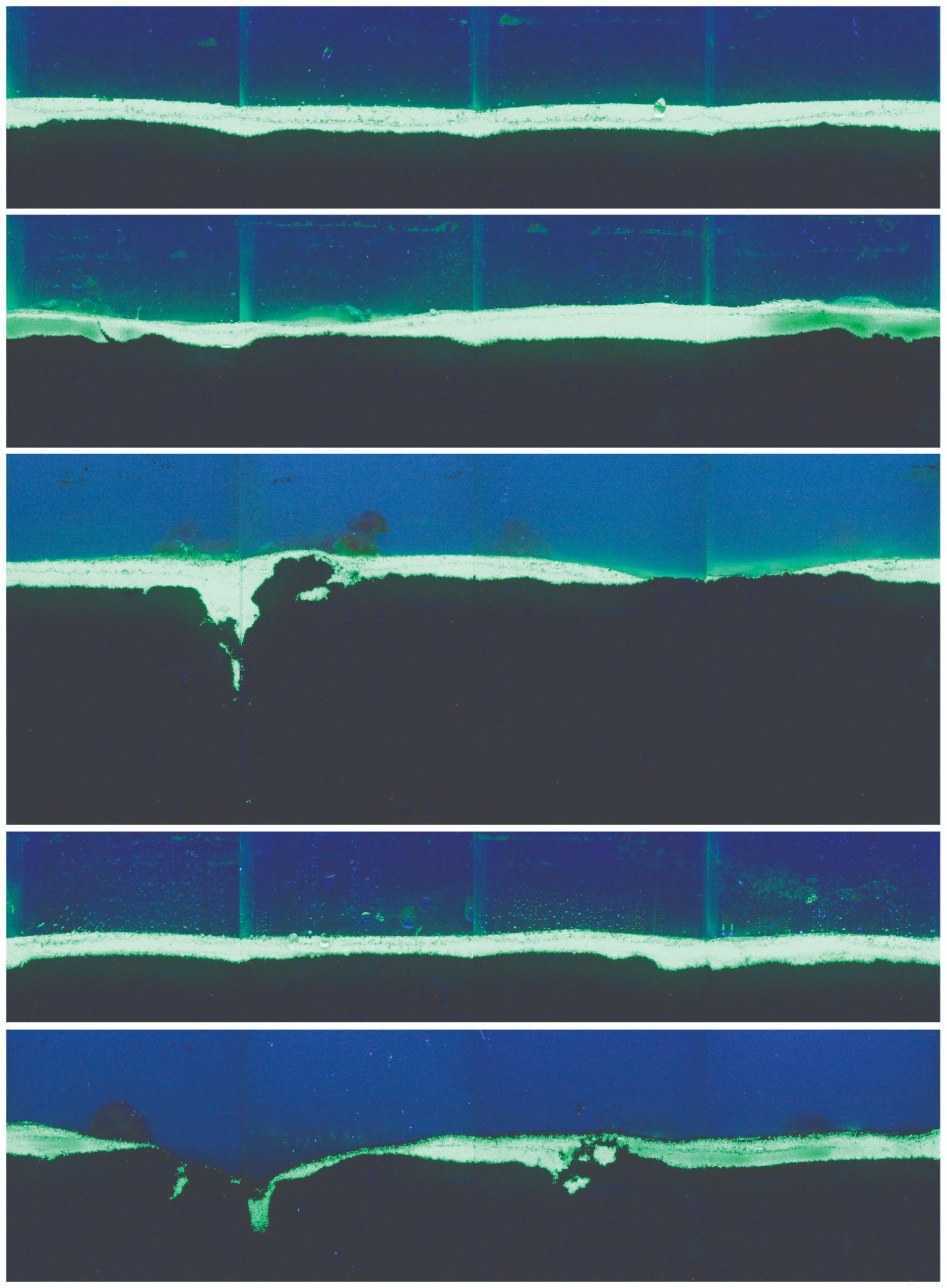

**Figure S14. f-SPI Images for *Terebellides stroemii* in 6 x 6 cm cores.** The five images show the five replicate cores.

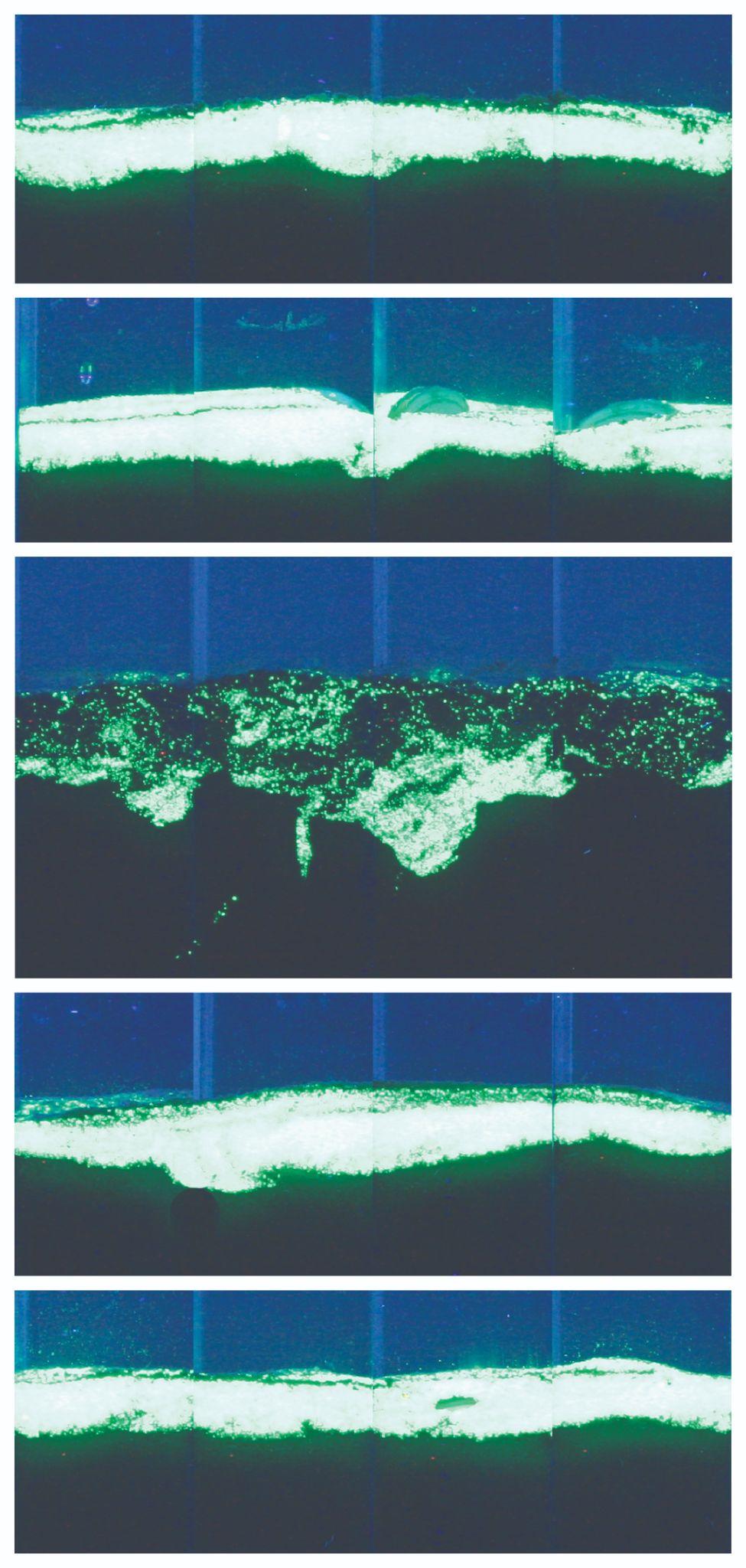

**Figure 15. f-SPI Images for *Abra nitida* in 2.2 x 2.2 cm cores.** The five images show the five replicate cores.

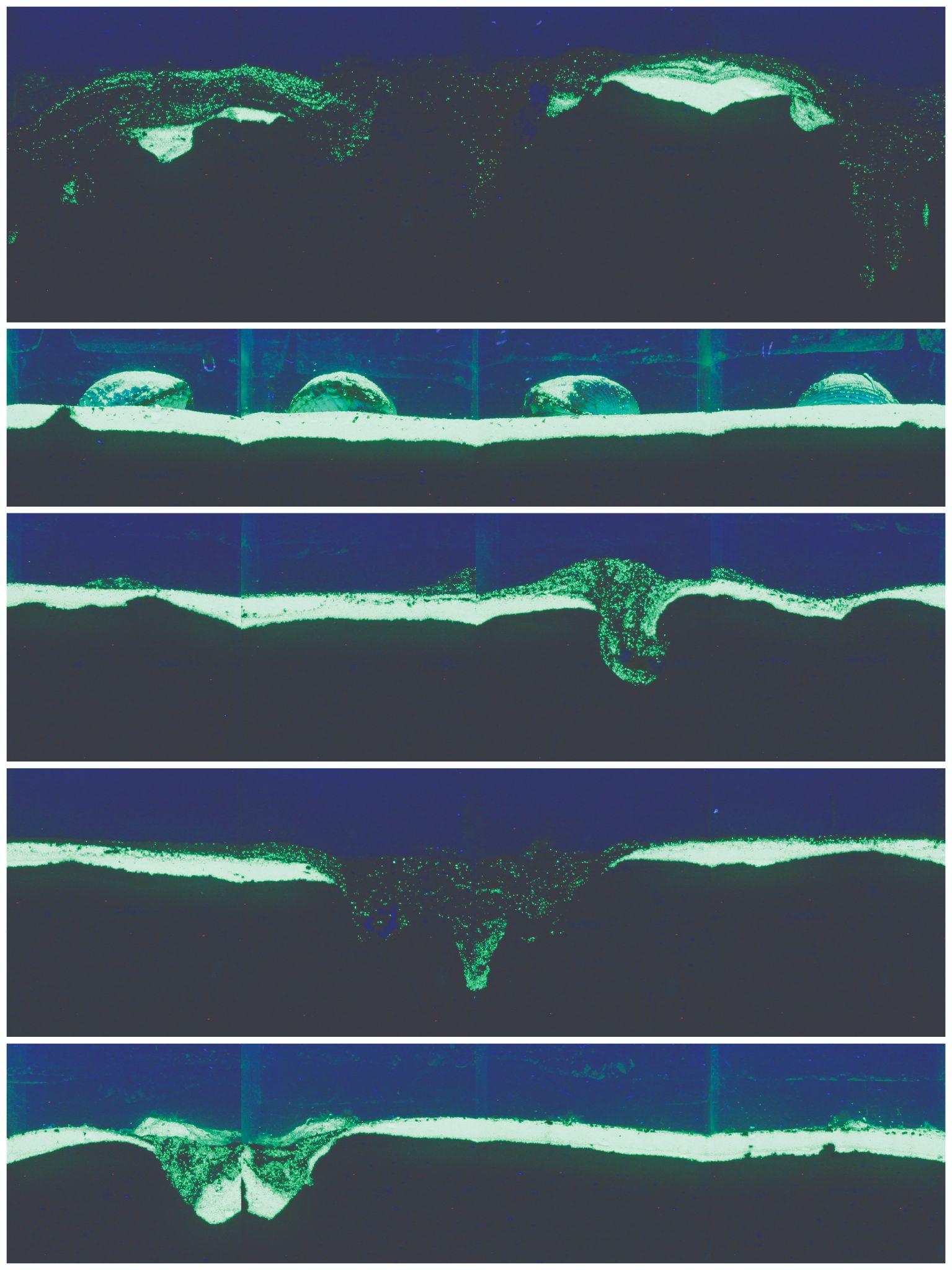

**Figure S16. f-SPI Images for *Cerastoderma edule* in 6 x 6 cm cores.** The five images show the five replicate cores.

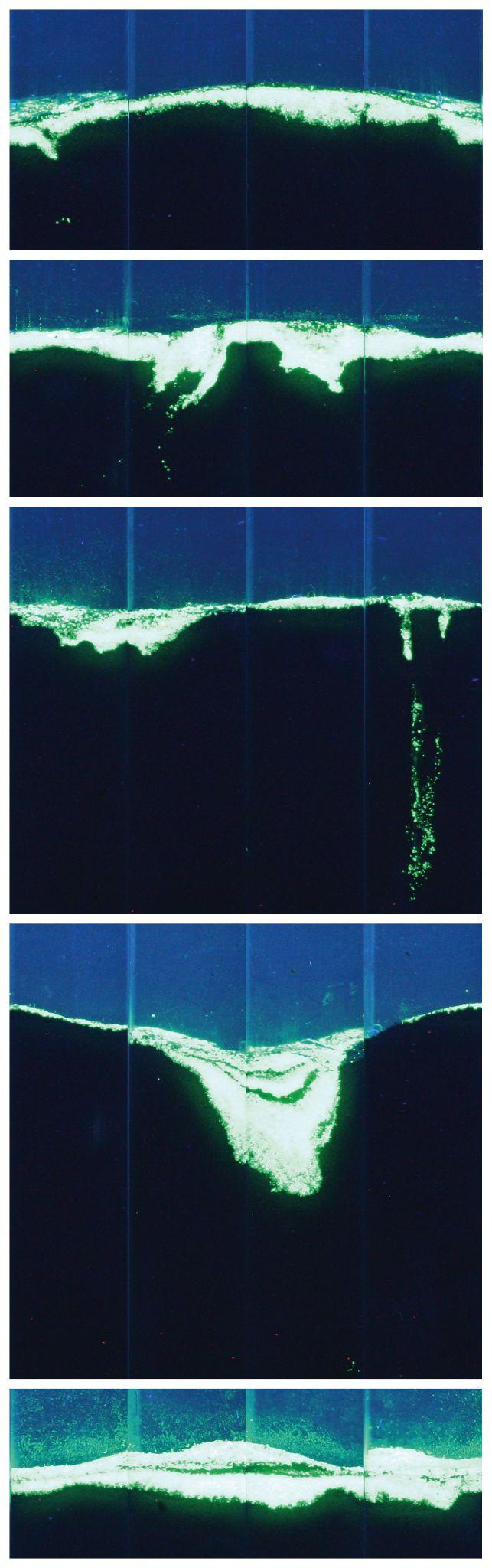

**Figure S17. f-SPI Images for *Mysia undata* in 2.2 x 2.2 cm cores.** The five images show the five replicate cores.

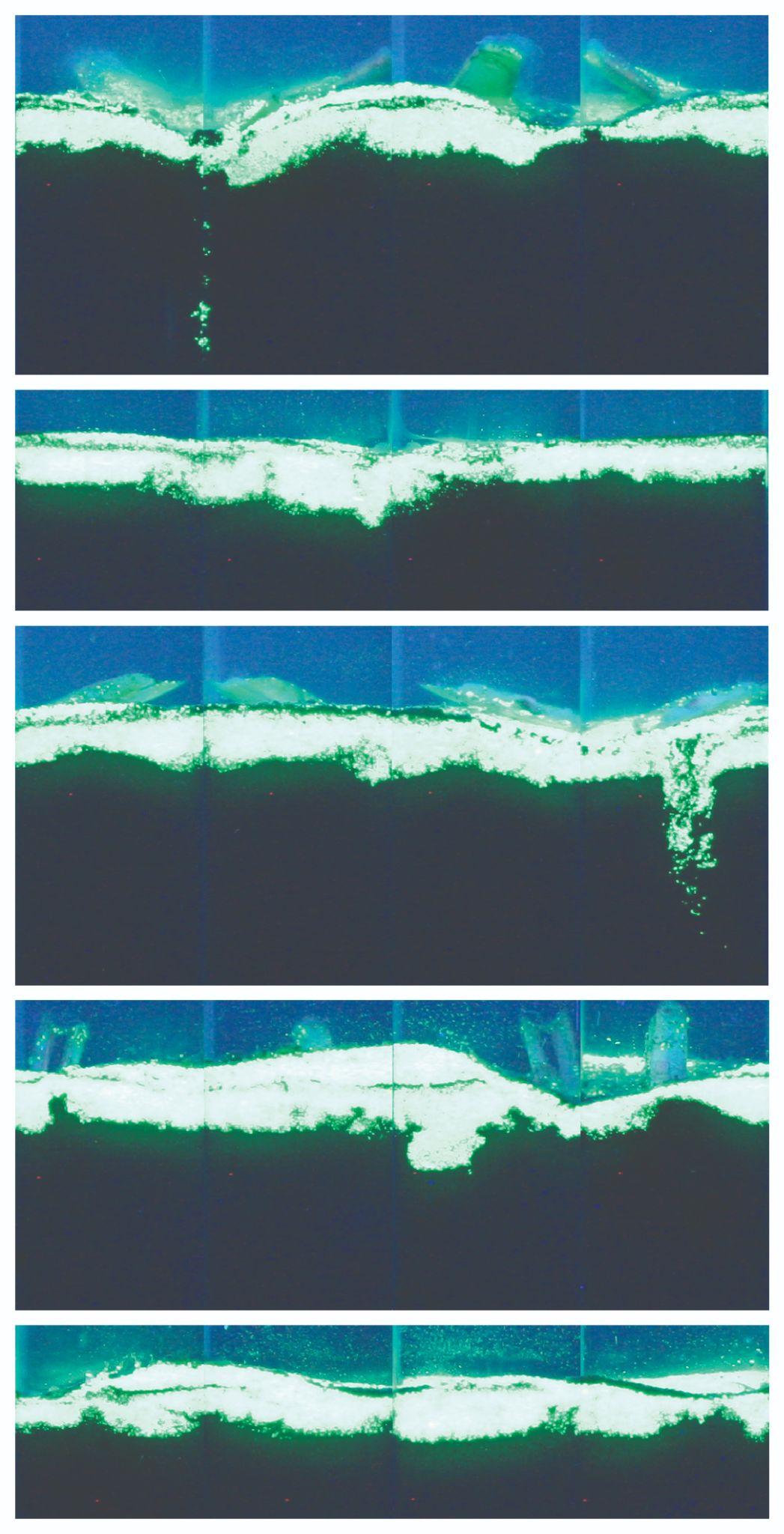

**Figure S18. f-SPI Images for *Phaxas pellucidus* in 2.2 x 2.2 cm cores.** The five images show the five replicate cores.

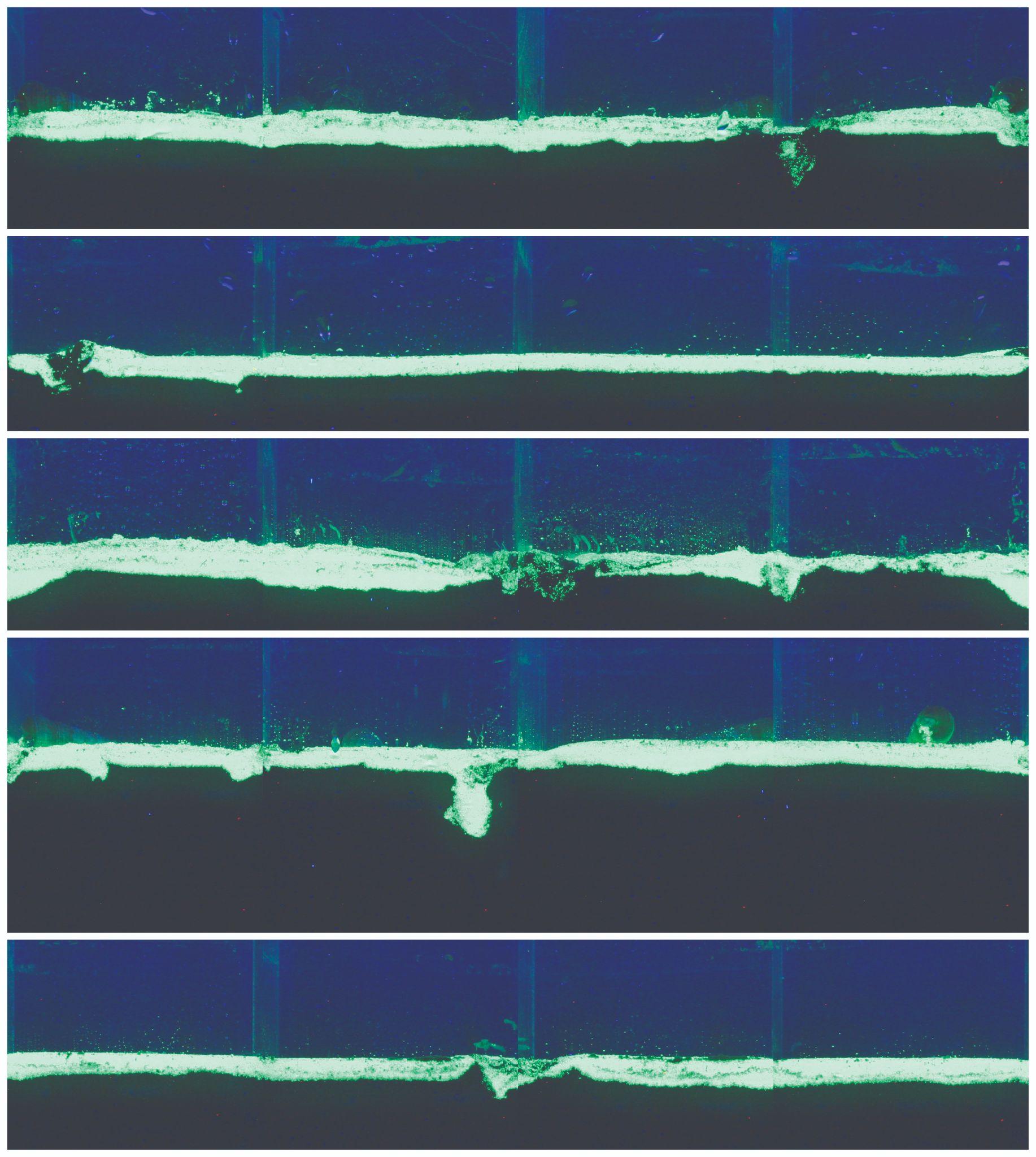

**Figure S19. f-SPI Images for *Turritellinella tricarinata* in 6 x 6 cm cores.** The five images show the five replicate cores.

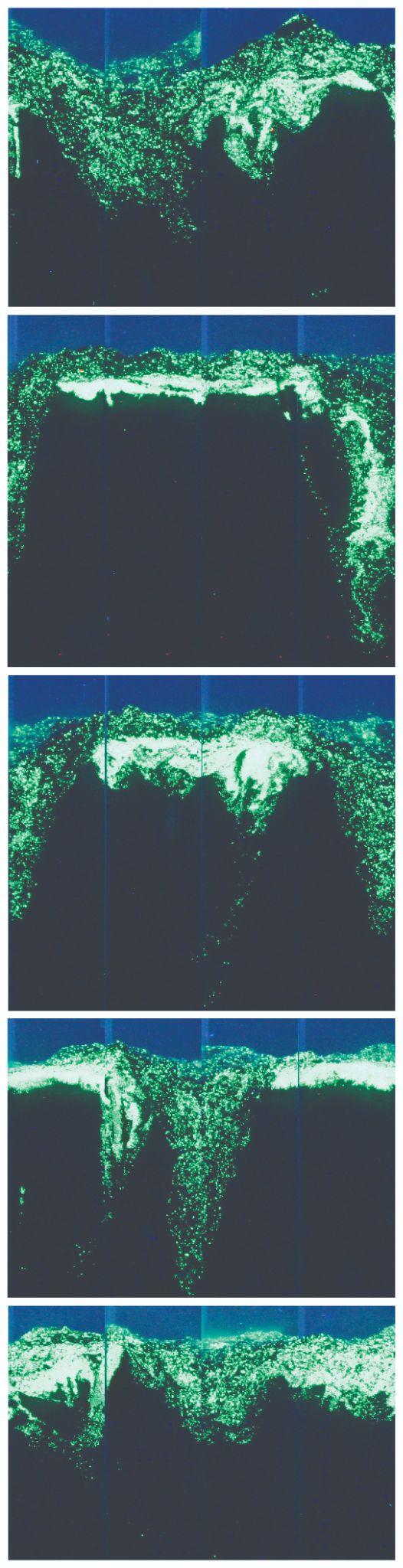

**Figure S20. f-SPI Images for *Amphiura chiajei* in 2.2 x 2.2 cm cores.** The five images show the five replicate cores.

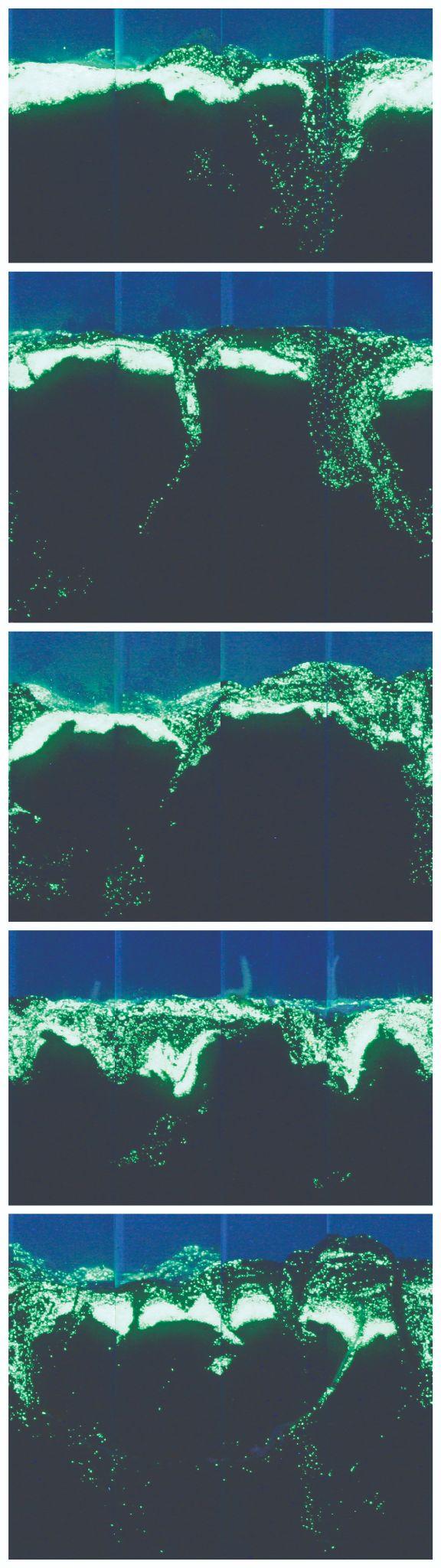

**Figure S21. f-SPI Images for *Amphiura filiformis* in 2.2 x 2.2 cm cores.** The five images show the five replicate cores.

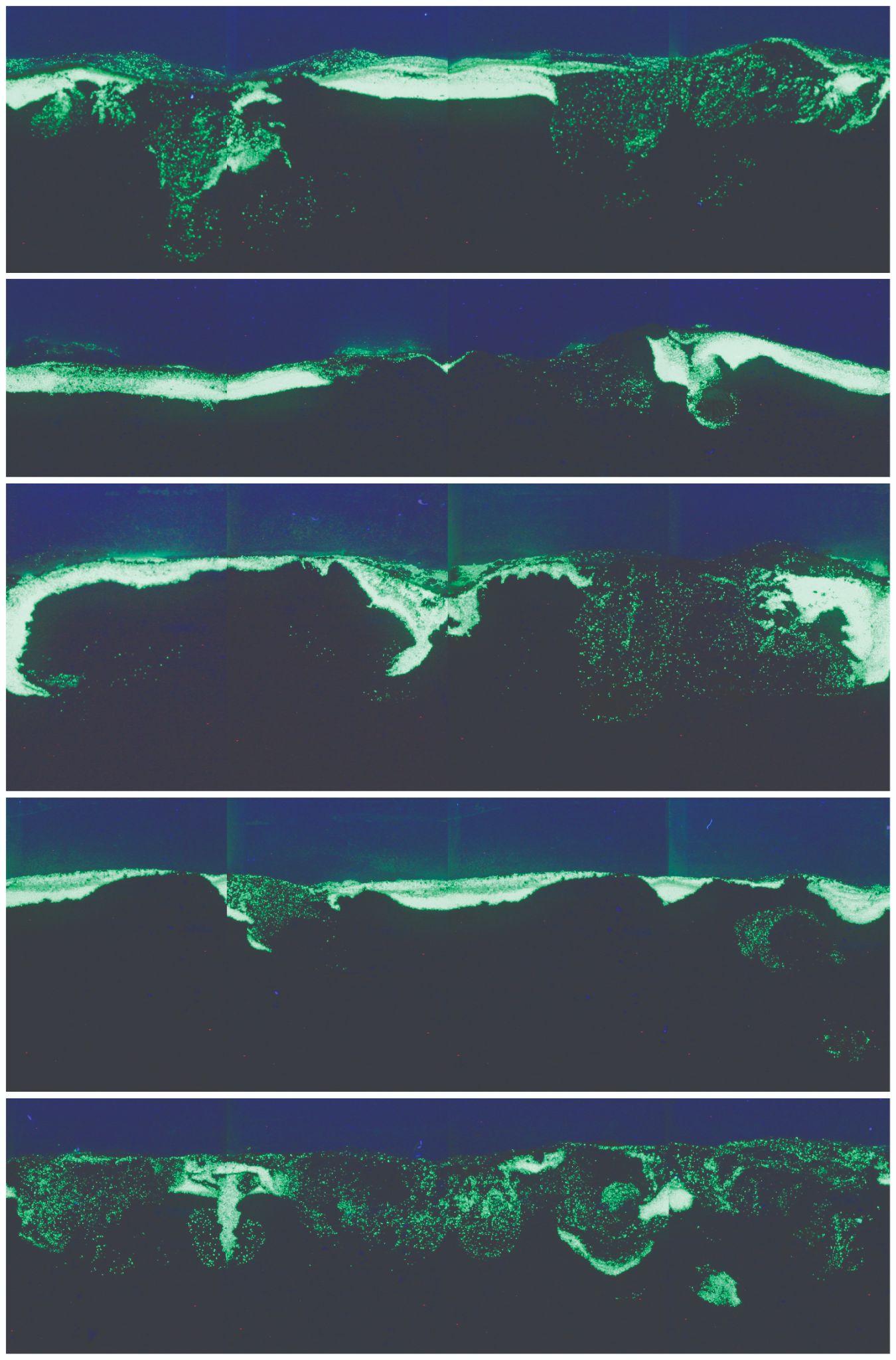

**Figure S22. f-SPI Images for *Echinocardium cordatum* in 6 x 6 cm cores.** The five images show the five replicate cores.

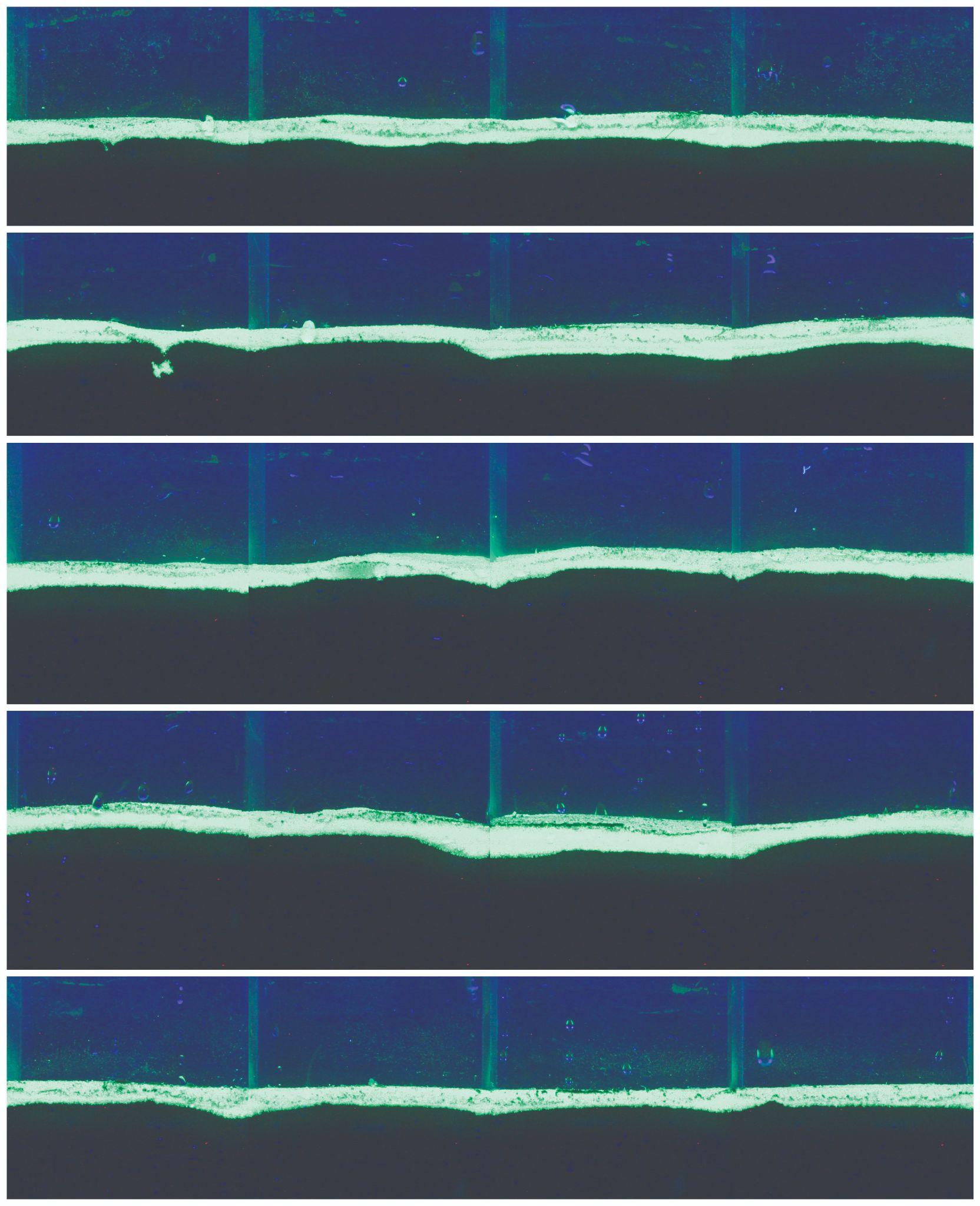

**Figure S23. f-SPI Images for *Paraleptopentacta elongata* in 6 x 6 cm cores.** The five images show the five replicate cores.

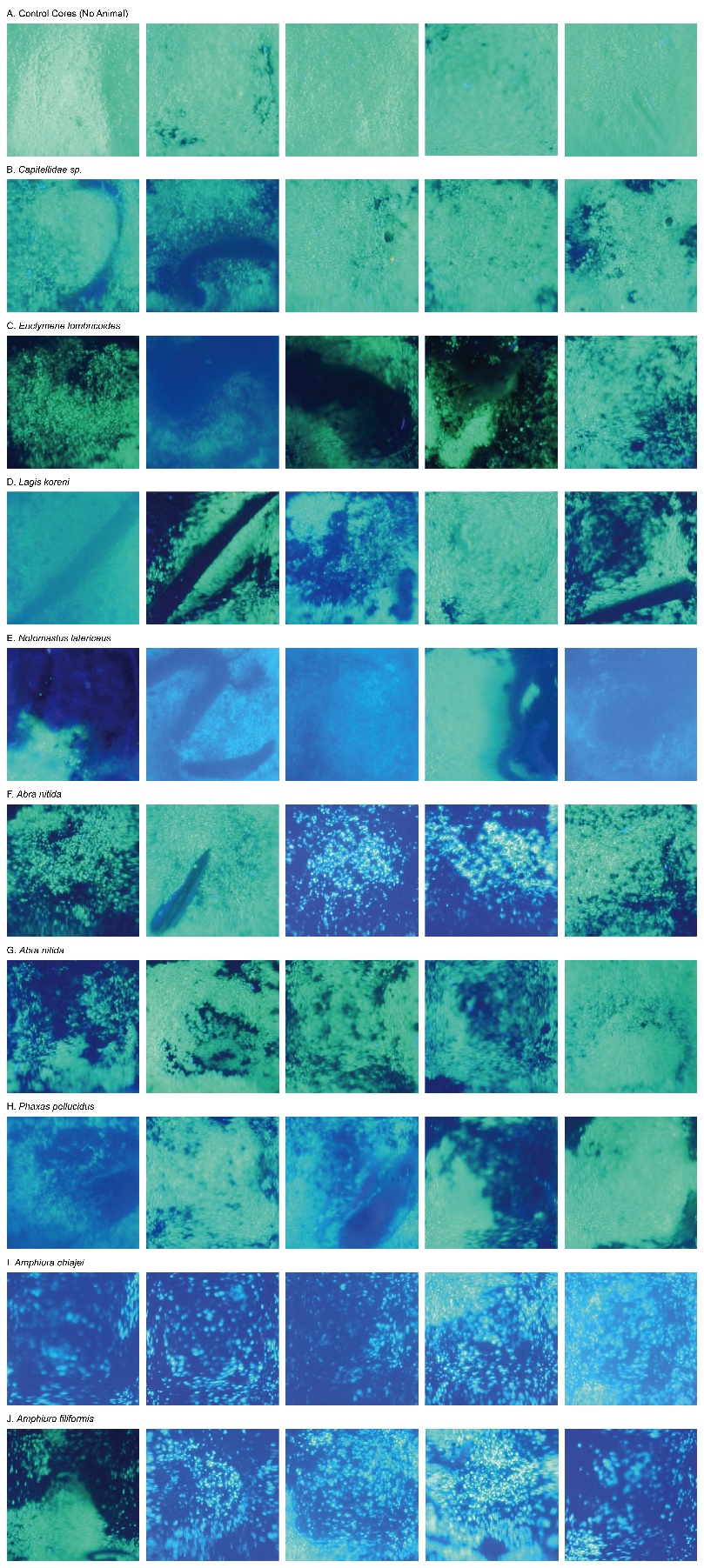

**Figure S24. Surface Sediment Imaging (SSI) Images for all species in the 2.2 x 2.2 cm cores.** The five images per species show the five replicate cores.

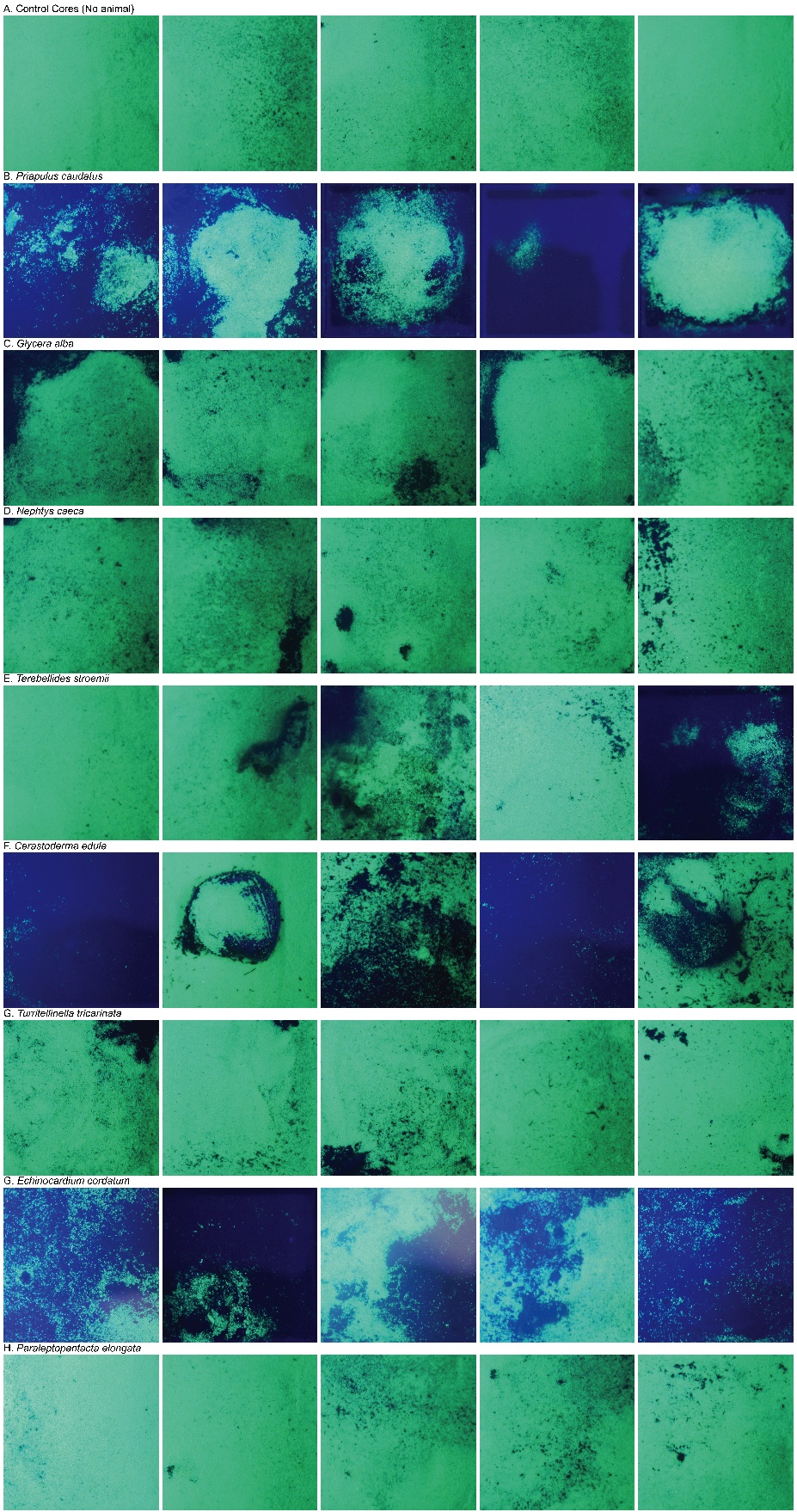

**Figure S25. Surface Sediment Imaging (SSI) Images for all species in the 6 x 6 cm cores.** The five images per species show the five replicate cores.

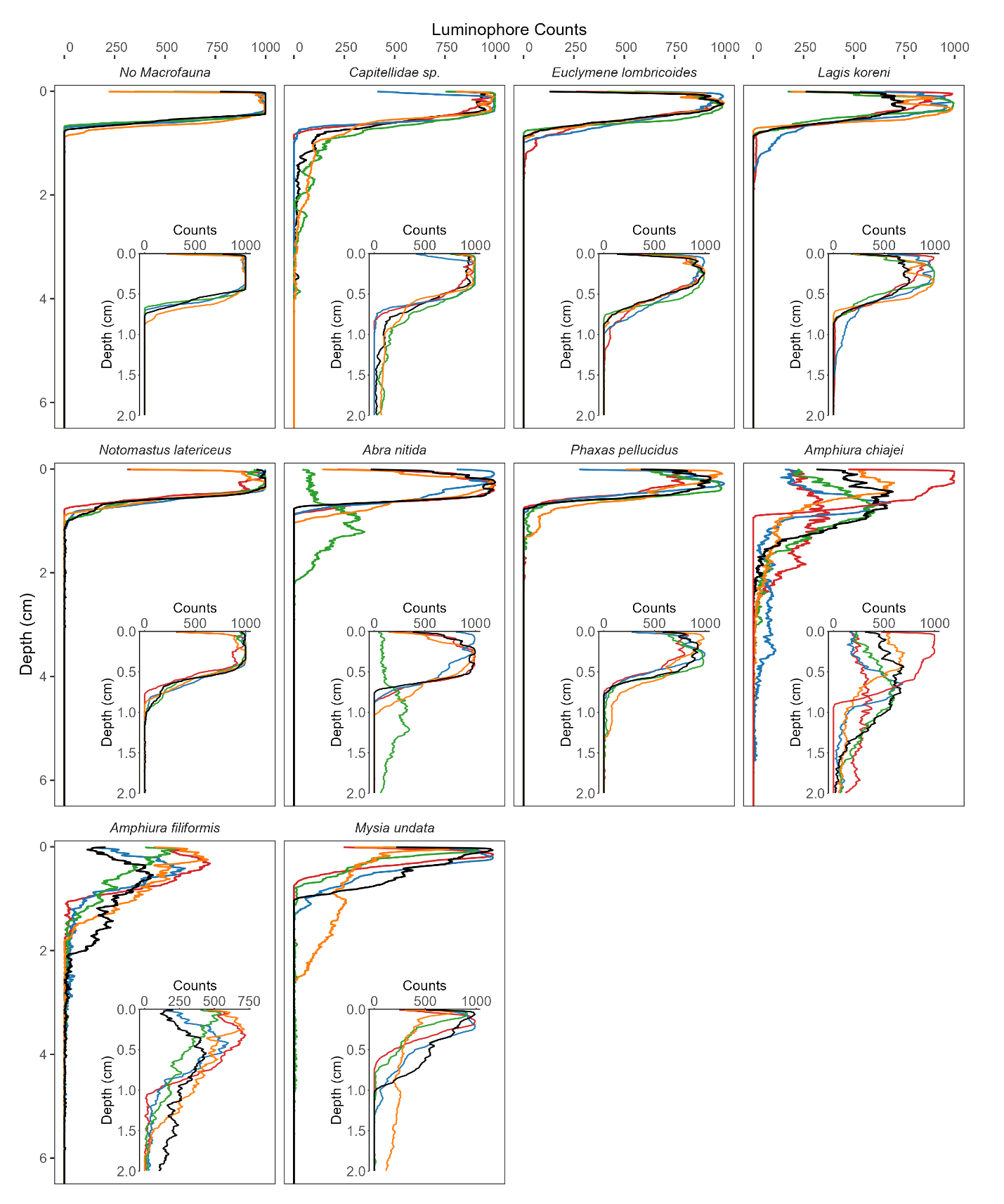

**Figure S26. Sediment particle reworking profiles (n=5) for all species in the 2.2 x 2.2 cm cores.** Insets show detail of main figure, colours represent individual replicates per species

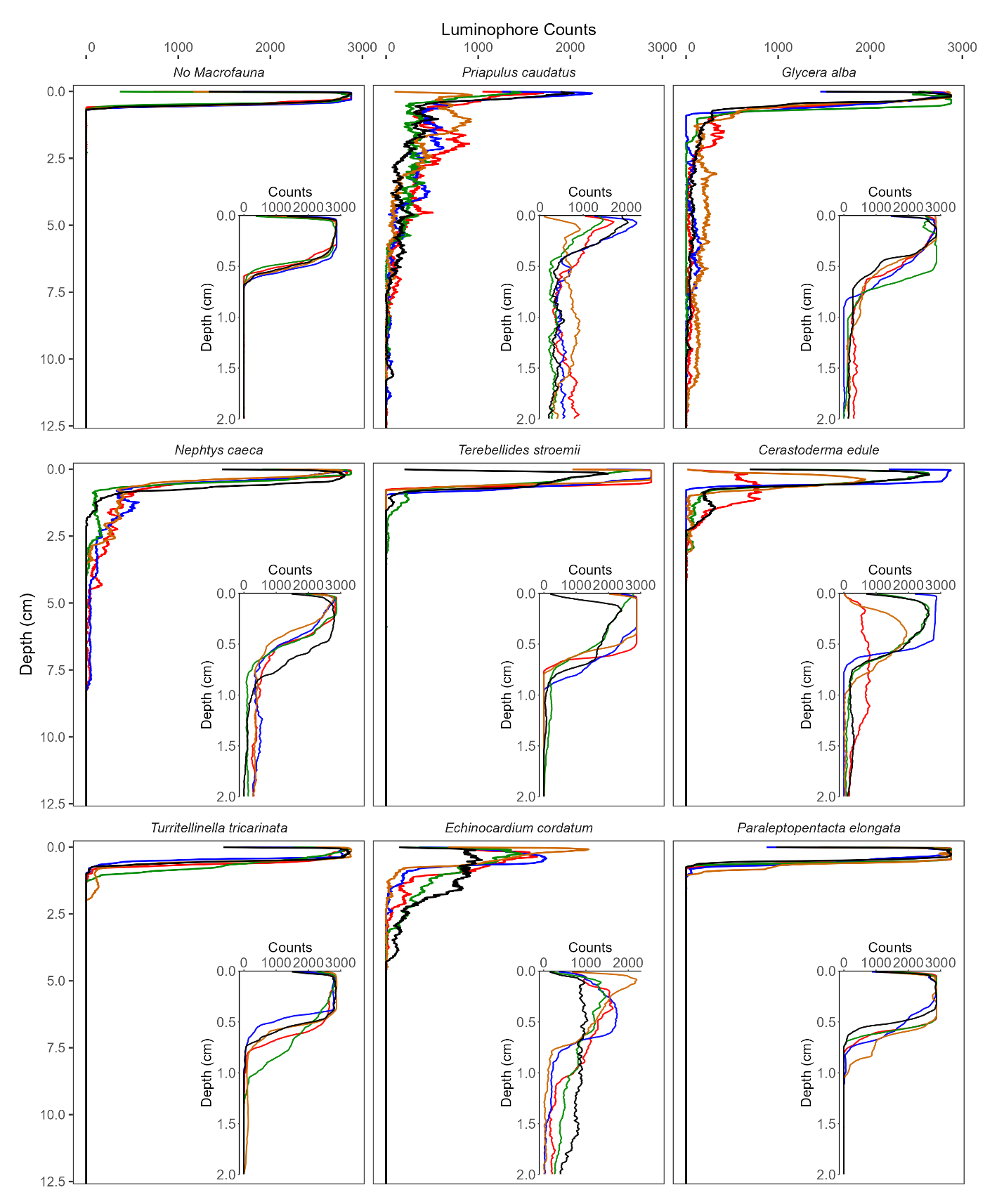

**Figure S27. Sediment particle reworking profiles (n=5) for all species in the 6 x 6 cm cores.** Insets show detail of main figure, colours represent individual replicates per species.
